# The MuvB Complex Binds and Stabilizes Nucleosomes Downstream of the Transcription Start Site of Cell-Cycle Dependent Genes

**DOI:** 10.1101/2021.06.29.450381

**Authors:** Anushweta Asthana, Parameshwaran Ramanan, Alexander Hirschi, Keelan Z. Guiley, Tilini U. Wijeratne, Robert Shelansky, Michael J. Doody, Haritha Narasimhan, Hinrich Boeger, Sarvind Tripathi, Gerd A. Müller, Seth M. Rubin

## Abstract

The chromatin architecture in promoters is thought to regulate gene expression, but it remains uncertain how most transcription factors (TFs) impact nucleosome position. The MuvB TF complex regulates cell-cycle dependent gene-expression and is critical for differentiation and proliferation during development and cancer. MuvB can both positively and negatively regulate expression, but the structure of MuvB and its biochemical function are poorly understood. Here we determine the overall architecture of MuvB assembly and the crystal structure of a subcomplex critical for MuvB function in gene repression. We find that the MuvB subunits LIN9 and LIN37 function as scaffolding proteins that arrange the other subunits LIN52, LIN54 and RBAP48 for TF, DNA, and histone binding, respectively. Biochemical and structural data demonstrate that MuvB binds nucleosomes through an interface that is distinct from LIN54-DNA consensus site recognition and that MuvB increases nucleosome occupancy in a reconstituted promoter. We find in arrested cells that MuvB primarily associates with a tightly positioned +1 nucleosome near the transcription start site (TSS) of MuvB-regulated genes. These results support a model that MuvB binds and stabilizes nucleosomes just downstream of the TSS on its target promoters to repress gene-expression.

## Introduction

Chromatin architecture and the position of nucleosomes influence DNA-mediated processes including the transcription of genes (Lai and Pugh, 2017). Transcription by RNA polymerase results in significant changes to nucleosome positioning as the basal transcription machinery must over-come the energetic barriers presented by the placement of nucleosomes along promoters and the gene body (Kujirai and Kurumizaka, 2020; Lorch et al., 1987; Teves et al., 2014). RNA polymerase with the aid of elongation factors and histone chaperones can bind and evict octamer proteins or reposition nucleosomes present in the gene body to access the gene for transcription. In addition, chromatin remodelers and histone modifying enzymes are thought to facilitate or inhibit transcription by arranging or displacing nucleosomes near transcription start sites, by altering the packing of nucleosomes, and by modulating the affinity of histone proteins for the DNA backbone. Less is known about how nucleosomal architecture is influenced by the activity of transcription factors (TFs). While recent evidence shows that pioneer TFs can bind target DNA sites within the nucleosome wrap and recruit remodelers to alter chromatin architecture, other TFs compete with nucleosomes for access to their DNA consensus sequence (Michael et al., 2020; Soufi et al., 2015; Zhu et al., 2018). A thorough molecular description of how many regulatory TFs cooperate and engage with nucleosomes to modulate gene-expression remains elusive.

The MuvB TF complex binds to target gene promoters and regulates a large set of cell-cycle genes. MuvB temporally coordinates the expression of genes necessary for DNA synthesis, centromere construction, mitotic division, and cell-cycle exit (Dynlacht, 1997; Fischer and Muller, 2017; Sadasivam and DeCaprio, 2013). In mammals, cell-cycle dependent gene expression occurs primarily in two waves of transcription which take place around the G1/S and G2/M transitions and depend on the activity of MuvB and the other TFs E2F, B-Myb, and FoxM1 (Bar-Joseph et al., 2008; Fischer et al., 2016; Grant et al., 2013; Liu et al., 2017; Whitfield et al., 2002). These TFs and their regulators are commonly deregulated in cancer (Kent and Leone, 2019; Musa et al., 2017; Myatt and Lam, 2007).

The MuvB complex, evolutionarily conserved throughout animals, plays a key role in development and differentiation and is considered a master regulator of cell-cycle dependent gene expression programs (Fischer and Muller, 2017; Harrison et al., 2006; Korenjak et al., 2004; Lewis et al., 2004; Litovchick et al., 2007; Sadasivam and DeCaprio, 2013; Schmit et al., 2007). During quiescence and in early G1, MuvB binds to the retinoblastoma protein (Rb) paralogs p130 or p107 (p130/p107) and E2F4-DP. This complex, known as DREAM, represses S phase genes and late cell-cycle genes (Litovchick et al., 2007; Mages et al., 2017; Schade et al., 2019b). Upon entry into the cell cycle, cyclin dependent kinases along with their cyclin partners phosphorylate and release p130/p107 from the MuvB core, disassembling DREAM but keeping the core MuvB intact (Guiley et al., 2015; Litovchick et al., 2007; Sandoval et al., 2009; Schade et al., 2019b). During S phase, the MuvB core binds to the onco-protein B-Myb and forms the Myb-MuvB (MMB) complex, which in concert with FoxM1 functions as a transcriptional activator of G2/M genes (Litovchick et al., 2007; Sadasivam et al., 2012; Schmit et al., 2007). While the cellular imbalance of activating and repressive MuvB complexes is associated with several cancers (Iness et al., 2019; Kim et al., 2021), the molecular details of MuvB assembly and function are poorly understood.

The core MuvB complex is composed of the five proteins LIN9, LIN37, LIN52, LIN54, and RBAP48. MuvB is localized to its target cell-cycle genes through LIN54, which binds target promoters directly at a consensus DNA sequence (Marceau et al., 2016; Muller et al., 2012; Schmit et al., 2009). The short sequence motif, known as the cell-cycle genes homology region (CHR), is found in close proximity to the transcription start site (TSS) and is often located four nucleotides downstream of a truncated E2F binding site, known as the cell cycle-dependent element (CDE) (Muller et al., 2014). LIN52 is a transcription factor adaptor protein that recruits either B-Myb or p130/p107, depending on cell-cycle phase (Guiley et al., 2018; Litovchick et al., 2011). RBAP48 is a histone binding chaperone protein that is found in several complexes that interact with chromatin, including CAF-1, NuRD, PRC2, and SIN3-HDAC (Schuettengruber et al., 2007; Verreault et al., 1996; Zhang et al., 1997; Zhang et al., 1999). In mammals, RBAP48 has a highly similar (89% sequence identity) paralog named RBAP46, which has not been identified in complexes with MuvB components (Litovchick et al., 2007). Both proteins are found in chromatin remodeler complexes and sometimes together. Less is known regarding the structure and biochemical function of LIN9 and LIN37, although a LIN9 sequence near its C-terminus co-folds with LIN52 to create the B-Myb binding site (Guiley et al., 2018).

Genetic evidence suggests that MuvB core proteins are essential in regulating cell-cycle dependent gene expression. In flies and worms, knockout of MuvB components contributes to inappropriate derepression of developmental gene programs (Harrison et al., 2006; Korenjak et al., 2004; Lewis et al., 2004). In mammals, LIN9 is essential for the expression of G2/M genes; loss of LIN9 causes mitotic defects and is embryonically lethal in mice (Osterloh et al., 2007; Reichert et al., 2010). On the other hand, knockdown of LIN9 in cell culture results in compromised repression of DREAM target genes upon induced cell-cycle exit (Litovchick et al., 2007). Knockout of the MuvB subunit LIN37 results in loss of MuvB-mediated gene repression in G0 and G1, but knockout does not lead to any observable changes in Myb-MuvB (MMB) gene expression in G2/M (Mages et al., 2017). Similarly, RNAi depletion of the *Drosophila* ortholog of RBAP48 specifically results in a derepression of dE2F2 target genes but does not result in defects in proliferation or gene expression (Taylor-Harding et al., 2004). These findings implicate MuvB core subunits in both positively and negatively modulating gene-expression, yet the biochemical mechanism behind their function remains unknown.

Here we investigated how MuvB represses gene expression, with emphasis on characterizing the structure and function of LIN9, LIN37, and RBAP48. We demonstrate that LIN9 and LIN37 together form an essential scaffold that holds together the core complex, and we determined a crystal structure that reveals how they together recruit RBAP48. We show that through RBAP48, MuvB binds directly to nucleosomes, either by interacting with H3 tails or the core particle. Using single-molecule electron microscopy, we found that MuvB increases nucleosome occupancy in a reconstituted cell-cycle gene promoter. These data indicate that MuvB associates with and stabilizes nucleosomes in the absence of other factors. Finally, we implemented a protocol that applies micrococcal nuclease digestion of chromatin and co-precipitation (MNase-ChIP) to study interactions of MuvB with nucleosomes in HCT116 cells. Our results support a model that MuvB binds to nucleosomes near the transcription start sites of target genes and stabilizes nucleosomes to repress cell-cycle dependent gene expression.

## Results

### LIN9 and LIN37 are together required for assembly of MuvB

Beyond the role of LIN9 in binding B-Myb, the structure and biochemical function of LIN9 and LIN37 have not been previously characterized. Human LIN37 is a conserved 246 amino acid protein that has no homology to any known structures. Sequence analysis suggests the presence of several short, structured regions (1-43, 95-126, 203-246) that are interspersed with sequences that are likely disordered (Figure 1A). The highly conserved segment 95-126, which we call the CRAW domain for the presence of a CRAW amino acid sequence, is necessary for LIN37 assembly into MuvB and for its activity in gene repression (Mages et al., 2017). Human LIN9 contains 538 amino acids, and beyond the presence of a Tudor domain, it also exhibits no homology to known structures (Figure 1A). The N-terminal ∼95 amino acids of LIN9 are poorly conserved and have no predicted structure. The next segment from 94-278 (previously called the domain in Rb-related pathway or DIRP; Pfam 06584) contains a Tudor domain and is conserved between MuvB and the related tMAC complex (White-Cooper et al., 2000). The helical segment between 349-466 forms the Myb-binding domain (MBD) together with LIN52 (Guiley et al., 2018), while the C-terminus (residues 473-538) also has predicted helical structure.

**Figure 1:**
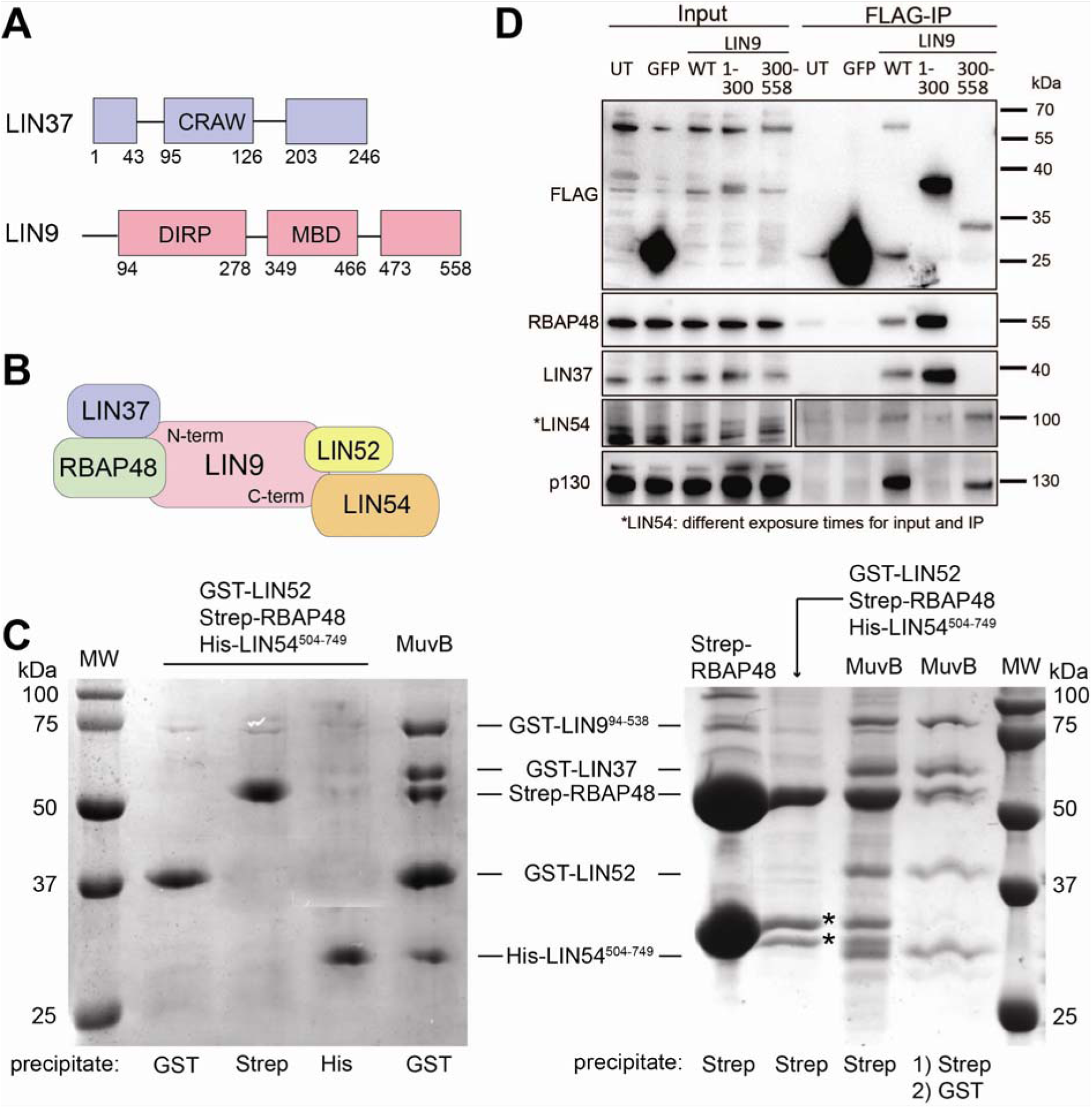
LIN9 and LIN37 scaffold the MuvB complex. (A) Domain architecture of LIN9 and LIN37, with regions of predicted and validated structure shown as blocks. The conserved LIN37 CRAW domain and LIN9 DIRP domain structures are determined here. MBD is the Myb-binding domain. (B) Schematic model for subunit interactions within MuvB. (C) The indicated tagged subunits or MuvB complex (GST-LIN52, Strep-RBAP48, His-LIN54^504-749^, GST-LIN37, and GST-LIN9^92-538^) were expressed in Sf9 cells and extracts were precipitated with resin capturing the indicated tag. Proteins were visualized with coomassie staining. *Indicates impurities or degradation observed in some RBAP48 expressions. These bands are not pulled out from the tandem purification. (D) HCT116 cells were transfected with plasmids encoding the indicated FLAG-tagged protein. FLAG-tagged proteins were precipitated from extracts using anti-FLAG antibody and visualized with anti-FLAG immunoblotting and immunoblotting with antibodies that recognize RBAP48, LIN37, LIN54, and p130.

Considering previous observations that LIN9 binds directly to multiple core MuvB and MuvB-interacting proteins (Guiley et al., 2018; Sandoval et al., 2009; Schmit et al., 2007) and that LIN9 knockdown results in DREAM complex assembly defects in T98G cells (Litovchick et al., 2007), we hypothesized that LIN9 is a scaffold onto which the other proteins assemble (Figure 1B). To probe MuvB complex assembly in a reconstituted system, we performed co-precipitation experiments by over-expressing proteins with different affinity tags in Sf9 insect cells (Figure 1C). We expressed full-length RBAP48, LIN52, and LIN37 and the relatively conserved and structured regions of LIN9 (residues 92-538, called LIN9^92-538^) and LIN54 (residues 504-749, LIN54^504-749^). When the three MuvB components RBAP48, LIN52, and LIN54 were co-expressed, we did not see co-precipitation (Figure 1C). In contrast, we were able to reconstitute the MuvB complex when all five core components were co-transfected, as demonstrated by performing successive precipitations of different affinity tags (Figure 1C). In our baculovirus system, we were unable to express LIN9 in the absence of LIN37, so we could not test whether LIN9 alone is required in our reconstitution. However, it has previously been reported that DREAM and MuvB complexes are able to assemble in the absence of LIN37 (Harrison et al., 2006; Mages et al., 2017; Uxa et al., 2019). Taken together, these results suggest that the LIN9 subunit of MuvB coordinates RBAP48, LIN52 and LIN54 to assemble the complex.

To further probe how LIN9 interactions with the other MuvB subunits organize the overall architecture of the complex, we expressed Flag-tagged LIN9 constructs in HCT116 cells and analyzed binding by co-immunoprecipitation (Figure 1D). We observed differences in the interactions made by LIN9^1-300^, which contains the DIRP domain, and the interactions made by LIN9^300-558,^, which contains the Myb-binding domain and C-terminus. Only LIN9^300-558^ co-precipitated p130. This observation is consistent with the known direct association of LIN9^MBD^ with LIN52 and the direct association of the LIN52 N-terminus with p130 (Guiley et al., 2018; Guiley et al., 2015). LIN9^300-558^ also associates with LIN54, whereas LIN9^1-300^ does not immunoprecipitate LIN54 above background in our experiment. In contrast, only LIN9^1-300^ co-precipitated RBAP48 and LIN37. We conclude that the LIN9 N-terminus is necessary and sufficient for binding RBAP48 and LIN37, while the C-terminus binds LIN52 and LIN54 (Figure 1B). We found that co-expression of RBAP48 with the DIRP region of LIN9 (LIN9^94-274^) and the conserved CRAW domain of LIN37 (LIN37^92-130^) in Sf9 cells yielded a MuvB subcomplex that was stable through affinity purification and size-exclusion chromatography (Figure S1). We call this subcomplex MuvBN, as it contains sequences toward the N-termini of LIN9 and LIN37.

### Structure of LIN9-LIN37-RBAP48 subcomplex

#### Overall structure

We were able to crystalize the MuvBN subcomplex, and we determined the structure to 2.55 Å by molecular replacement using the known RBAP48 subunit structure as an initial model (PDB: 3GFC) (Xu and Min, 2011). The crystal structure contained one complex in the asymmetric unit, and we built the LIN9 and LIN37 fragments into the unmodeled electron density. The final refined MuvBN model contains one copy of each protein (Figure 2). As previously described, RBAP48 has a β-propeller domain fold, consisting of seven small β-sheets, along with a single N-terminal helix (Schmitges et al., 2011; Song et al., 2008; Xu and Min, 2011). The atomic structure of RBAP48 in MuvBN aligns well with other structures of the protein in other complexes with RMSDs ∼0.3-0.6 Å (Figure S2). The LIN9^94-274^ sequence is almost entirely visible in the electron density and contains six alpha helices and the Tudor domain. The helices are N-terminal to the Tudor domain and do not appear to form a globular structure. Instead, they wrap around and from extensive contacts with RBAP48, create a binding site for LIN37, and anchor the Tudor domain to the rest of the complex. The LIN37^92-130^ CRAW domain is also nearly all visible in the electron density. This continuous LIN37 sequence forms two small β-strands and a short α helix. Our recombinant LIN9 was unstable without co-expression of this highly conserved fragment of LIN37^92-130^, which interacts with both RBAP48 and LIN9 in the subcomplex.

**Figure 2:**
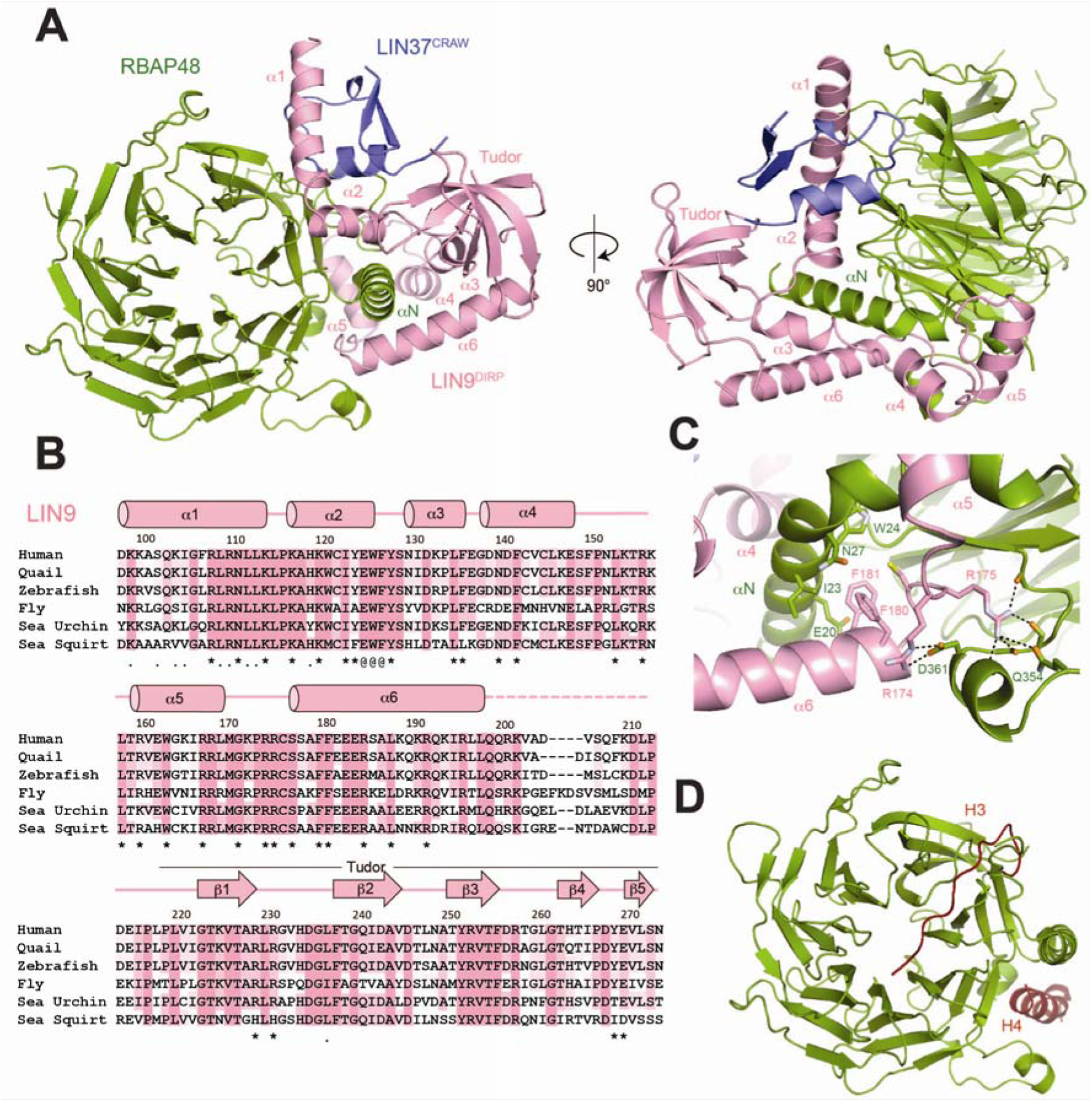
Structure of the MuvBN subcomplex. (A) Overall structural model. (B) Alignment of LIN9 sequences from *H. sapiens*, *C. japonica*, *D. rerio*, *D. melanogaster*, *S. purpuratus*, and *C. intestinalis*. The * marks residues that contact RBAP48, the . marks residues that contact LIN37, and the @ marks residues that contact both. (C) Close-up view of one interface between LIN9 and RBAP48. (D) Location of histone H3 and histone H4 peptide binding sites on RBAP48. When bound to LIN9, the H4 sites is blocked while the H3 site is mostly accessible. The model was generated from PDB IDs: 2YBA and 3CFV.

#### Structure of the LIN9-RBAP48 interface

LIN9 and RBAP48 associate across a broad interface focused around the N-terminal helix (αN) of RBAP48 and the adjacent side of the β-propeller domain (Figure 2). All six LIN9 helices contact RBAP48, and five of them (α2-α6) surround and make interactions with the RBAP48 αN helix (Figures 2A and 2B). Numerous hydrophobic, polar, and electrostatic contacts are observable between the proteins (Figure 2B), and we highlight a few specific examples here that are relevant for the mutagenesis experiments described below. For example, a cluster of two arginines (R174 and R175) and two phenylalanines (F180 and F181) in LIN9 anchor α6 and the preceding loop against the RBAP48 αN helix and so-called PP-loop, which is an insertion in the sixth propeller β-sheet (Figure 2C). The sidechains of R174 and R175 make a series of electrostatic interactions with side chain and main chain atoms in RBAP48 residues Q354, D358, P361, and G362, while F180 and F181 pack against RBAP48 residues I23 and W24.

Structures of RBAP48/RBAP46 bound with peptides depict how they are assembled into diverse complexes. A survey of known structures in complex with peptides reveals two common binding sites on the β-propeller domains (Figures 2D and S2). One site is across the face of the β-propeller and is found occupied by histone H3, Fog1, and PHF6. The second site is along the side of the propeller between αN and the PP loop; it is found occupied by histone H4, Mta1, and Suz12. In the MuvBN structure, the H4 site is bound by the sequence in LIN9 between α5 and α6. In contrast, the H3 site is for the most part accessible, although the α1 and α2 helices of LIN9 pack against the edge of the propeller where the H3 site-binding peptides exit the propeller face.

Several structures of RBAP48 in complex with one or more proteins or larger protein fragments have been previously determined. For example, RBAP48 is present in the polycomb complex PRC2 (Chen et al., 2018; Schuettengruber et al., 2007). As observed in MuvBN, RBAP48 is bound in these other complexes at multiple sites and on both sides of αN. One striking difference in how LIN9 and LIN37 bind RBAP48 compared to how proteins bind in other complexes is the extensive interactions with a glycine rich loop in RBAP48 (residues 88-115) (Figure 3A). This RBAP48 loop, which is an insertion between two strands in the first complete propeller blade, is disordered in almost all the structures with peptides and partially ordered when binding Mat1 or the polycomb complex protein Suz12 (Figure S2). In contrast, the interactions of the insertion loop with LIN9 and LIN37 are much more extensive, and the entire loop appears ordered in the MuvBN structure.

**Figure 3:**
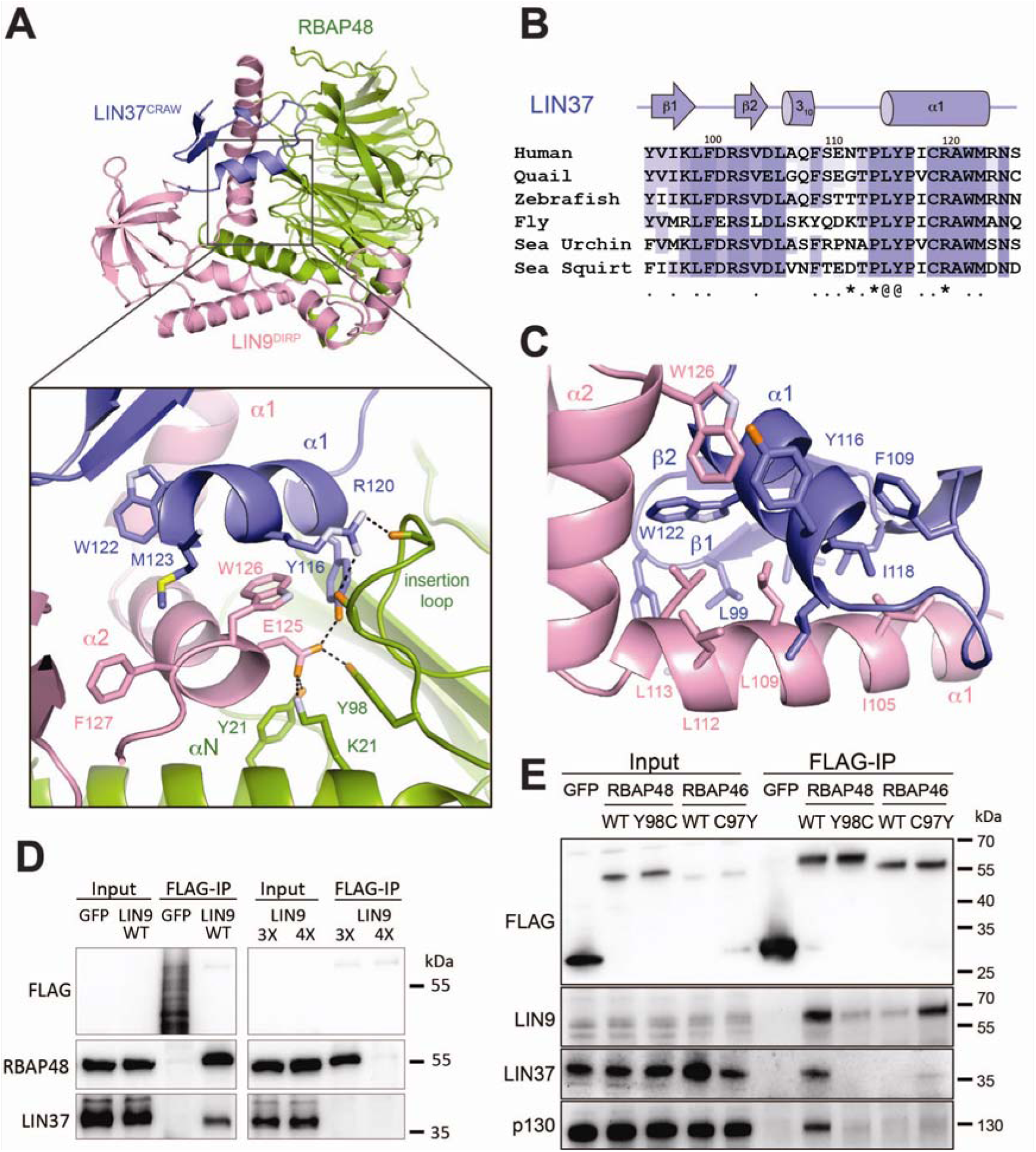
LIN37 CRAW domain binds both LIN9 and RBAP48. (A) Interactions of LIN9 and LIN37 with the RBAP48 insertion loop. (B) Alignment of LIN37 sequences from organisms as in Figure 2B. Residues that contact LIN9 (.), RBAP48 (*), and both LIN9 and RBAP48 (@) are indicated. (C) Close-up view of the LIN9-LIN37 interface. (D) The indicated FLAG-GFP control and FLAG-LIN9 WT and mutants were expressed by transient transfection in arrested HCT116 cells. Proteins were immunoprecipitated using anti-FLAG beads and the indicated proteins visualized by Western blot. The 3x LIN9 mutant is E125A/W126A/F127A and the 4x LIN9 mutant is R174A/R175A/F180A/F181A. (E) Same as (D) but expressing the indicated RBAP46 and RBAP48 WT and mutant proteins.

#### LIN9 Tudor domain has a non-canonical aromatic cage

LIN9 additionally contains a conserved Tudor domain that is visible in the subcomplex (residues 223-273). Tudor domains are protein interaction modules that are found in many chromatin-interacting proteins. In several cases, they recognize methylated lysines and arginines and function as readers of modified histones (Cai et al., 2013; Lu and Wang, 2013; Tripsianes et al., 2011). The Tudor structure is defined by five anti-parallel β strands that fold into a barrel. Target peptides are bound by an aromatic cage at one end of the barrel. The cage typically surrounds the modified basic sidechain and makes stabilizing π-cation interactions. We aligned the LIN9 Tudor domain with structures of the PHF1 (PDB: 2M0O, RMSD 1.0 Å) and the SMN (PDB: 4A4E, RMSD 0.9Å) Tudor domains in complex with their target peptides (Figure S3) (Cai et al., 2013; Tripsianes et al., 2011). The alignments suggest that the LIN9 cage contains fewer aromatics and is relatively inaccessible, as it makes an interaction with a loop that adjoins β3 and β4 at L261 (Figure S3). We note that we have not been able to detect binding of the LIN9 Tudor domain to several unmodified and modified histone peptides or modified lysine and arginine at high concentration. While we do not rule out the possibility that the LIN9 Tudor domain binds histones or other proteins, we conclude that the structural features of the cage that mediate the interactions of other Tudor domains are not obviously present in LIN9.

#### LIN37 structure and interface with LIN9 and RBAP48

Previous functional domain mapping studies demonstrated that two highly conserved sequences in LIN37 were critical for LIN37 binding to other DREAM components and for DREAM repression of cell-cycle gene expression (Mages et al., 2017). These sequences in LIN37 correspond with the CRAW domain of LIN37 that appears structured in our crystals of MuvBN, and they play a critical role in interacting with LIN9 and RBAP48. This observation firmly implicates the MuvBN subcomplex as the structural subunit of DREAM responsible for gene repression.

The small structured LIN37 CRAW domain is bound between LIN9 α1 and α2 (Figures 3A-3C). The two LIN9 helices form a V-shape that straddles one face of the LIN37 structure. Sidechains along one hydrophobic face of the LIN9 α1 helix (I104, L108, L111, and L112) are inserted into a groove formed by hydrophobic residues from all the LIN37 secondary structure elements (I97, L99, F100, V104, L106, F109, L115, I118, and W122). The LIN9 α2 helix binds the opposite face of the LIN37 helix from the LIN9 α1 helix with LIN9 W125 packing against the LIN37 backbone and interacting with LIN37 Y116. The LIN37 helix forms the primary interface between LIN37 and RBAP48 (Figure 3A). Y116 and R120, both of which are highly conserved among LIN37 orthologs, make several interactions with the glycine rich insertion loop in RBAP48. The nearby LIN9 α2 helix also contributes to this interface such that E124 from LIN9, Y116 from LIN37, and Y98 from RBAP48 all interact through a network of hydrogen bonds.

We tested the importance of several interface contacts observed in the structure on assembly of MuvB in HCT116 cells (Figure 3D). We expressed either FLAG-tagged WT LIN9 or two FLAG-tagged LIN9 mutants and performed anti-FLAG immunoprecipitation to assay association with other MuvB proteins. A triple mutant (E125A/W126A/F127A) that contains mutations in the LIN9 α1 helix (LIN9^3X^) failed to co-precipitate LIN37, whereas a quadruple mutant (R174A/R175A/F180A/F181A, LIN9^4X^) with mutations in LIN9 α6 and the preceding linker (Figure 2C) failed to co-precipitate both LIN37 and RBAP48. It is notable that LIN37 was lost in the LIN9^4X^ co-precipitation even though the mutated residues are not directly at the LIN37 interface. These results indicate that despite the extensive interface, RBAP48 association with LIN9 can be disrupted through a few key mutations. The results of the LIN9^4x^ mutant also suggest that LIN37 association with LIN9 is likely stabilized by the presence of RBAP48 in the complex.

Analysis of the interactions at the LIN9-LIN37-RBAP48 interface also reveal the structural mechanism for the specificity of RBAP48 in the MuvB complex. Previous analysis of MuvB components using mass spectrometry did not identify the presence of the RBAP48 paralog RBAP46 (Litovchick et al., 2007). In our co-immunoprecipitation experiments, we also did not observe association of RBAP46 with components of the complex (Figure 3E). The two human homologs are 89% identical, but notably RBAP46 contains a cysteine at position Y98 in RBAP48. In the MuvBN structure, Y98 is in the RBAP48 insertion loop and is involved in a network of hydrogen bonds at the interface with both LIN9 and LIN37 (Figure 3A). We found that while Flag-tagged wild-type mouse RBAP48 could co-precipitate MuvB components in HCT116 cells extracts, mouse RBAP48 with an RBAP46-mimicking Y98C mutation does not co-precipitate MuvB components (Figure 3E). Conversely, a mouse RBAP48-mimicking C97Y mutation in mouse RBAP46 results in some additional affinity, although we note association still appears weaker than with WT RBAP48. We conclude that the MuvB complex has specificity for RBAP48 and that this specificity arises through this unique insertion loop association with LIN9 and LIN37.

### MuvB binds histone H3 tails and reconstituted nucleosomes lacking a CHR site

The MuvB complex contains two domains that have potential histone binding properties: the Tudor domain of LIN9 and the β-propeller domain of RBAP48. We wanted to test whether these domains, within the context of MuvB, are able to engage with histone peptides as well as nucleosomes. We first tested whether our recombinant purified MuvB complexes bind histone peptides that are known to form complexes with RBAP48. We tested binding of both MuvB (Figure 4A) and the MuvBN subcomplex (Figure S4) to fluorescein labeled H3 (1-21) and H4 (21-41) peptides by fluorescence polarization. We found that both MuvB and MuvBN bound the H3 tail but that they did not bind the H4 peptide. This observation is consistent with the MuvBN structure, which shows that the H3 site in RBAP48 is accessible while the H4 site is occluded by LIN9 (Figure 2D). We found that MuvBN binds H3 peptide with similar but slightly weaker affinity as the full MuvB complex, suggesting that the MuvBN complex is sufficient to make the most significant contacts with the H3 peptide (Figure S4).

**Figure 4:**
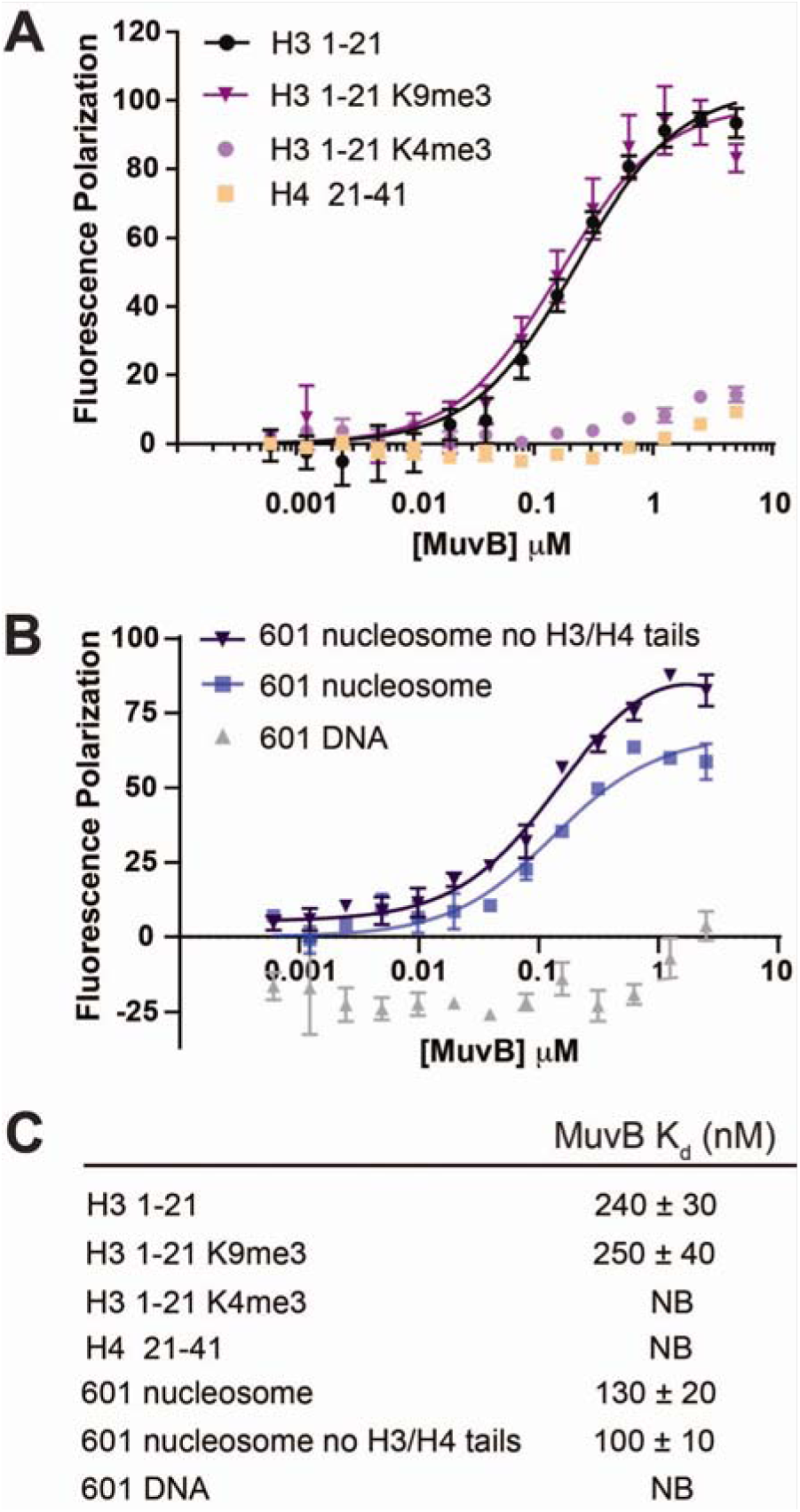
MuvB binds to histone peptides and nucleosomes. Fluoresence polarization measurements of association between recombinant MuvB and dye-labeled histone peptides. (A) or nucleosomes (B). FP is normalized to dye-labeled probe alone. (C) Average affinities from three replicates.

To probe whether post-translational modifications on H3 tails influence MuvB binding, we tested two H3 marks that are associated with active (H3K4me) or repressed (H3K9me3) transcriptional activity. We found that both MuvB bound H3 tails when methylated at K9 but failed to bind with H3 tails methylated at K4 (Figure 4A). This result is consistent with available structural data demonstrating that K4 methylation inhibits H3 tail binding to RBAP48 (Schmitges et al., 2011). We note that our observations contrast with experiments performed with purified *Drosophila* dREAM complex, which appeared to bind non-acetylated H4 peptides (Korenjak et al., 2004). However, the Drosophila complex contains additional histone-interacting proteins (L3MBT and an HDAC ortholog) not present in the mammalian complex.

We then asked whether MuvB could bind reconstituted nucleosomes and whether nucleosome binding was conferred by H3 tails alone. We reconstituted nucleosomes with full-length histones and the Widom 601 strong positioning sequence containing a fluorescein label. MuvB bound to these nucleosomes with slightly greater affinity than to the tails (Figures 4B and 4C). The 601 DNA sequence lacks a CHR sequence, and we found that MuvB did not bind fluorescein-labeled free 601 DNA, indicating that nucleosome association occurs independent of DNA consensus motif binding. We also observe MuvB interactions with reconstituted nucleosomes in a gel electromobility shift assay (Figure S4A). In the FP assay, we found that MuvB binds to the nucleosomes lacking histone H3 and H4 N-terminal tails (H3:39-136; H4:19-103) with a similar affinity compared to nucleosomes with tails (Figure 4B). This observation is consistent with a known association of RBAP48 with tailless histone H3-H4 dimers, although we detect here association in the context of a reconstituted nucleosome (Zhang et al., 2013). Our data indicate that MuvB can bind nucleosomes through the H3 tails but that H3-tail binding is not necessary for MuvB-nucleosome association. To rule out any potential binding of the LIN9 Tudor domain, we reconstituted a mutant MuvB complex harboring LIN9 Tudor aromatic cage mutations (L230A/F238A/F256A/H264A) and found this mutant engages with Widom nucleosomes similar to the wild-type complex in a gel shift assay (Figure S4). This result suggests that the LIN9 aromatic cage is not necessary for binding nucleosomes. Considering these results together, we propose that MuvB engages with H3 tails and/or the folded octamer to bind nucleosomes and that this association is primarily mediated by the MuvBN subcomplex including RBAP48.

### MuvB binds and stabilizes nucleosome occupancy on a reconstituted and chromatinized cell-cycle gene promoter

We previously analyzed late cell-cycle genes in available ENCODE data sets and found that DREAM target gene promoters show a higher nucleosome density within the few hundred bases downstream from the transcription start site relative to genes that lack a CHR site and to constitutively expressed genes (Marceau et al., 2016). Following our observation here that MuvB binds nucleosomes in the absence of additional factors, we tested whether MuvB directly increases nucleosome occupancy on cell-cycle gene promoters. We cloned and amplified a minimal promoter from the human *TTK* gene, which is a late cell-cycle gene that contains a single CHR and is regulated by MuvB (Muller et al., 2017). We folded a purified TTK-derived 461bp DNA fragment with recombinant histone octamer in the presence and absence of MuvB. An electromobility shift assay demonstrated that MuvB was able to associate with the chromatinized promoter (Figure 5A).

**Figure 5:**
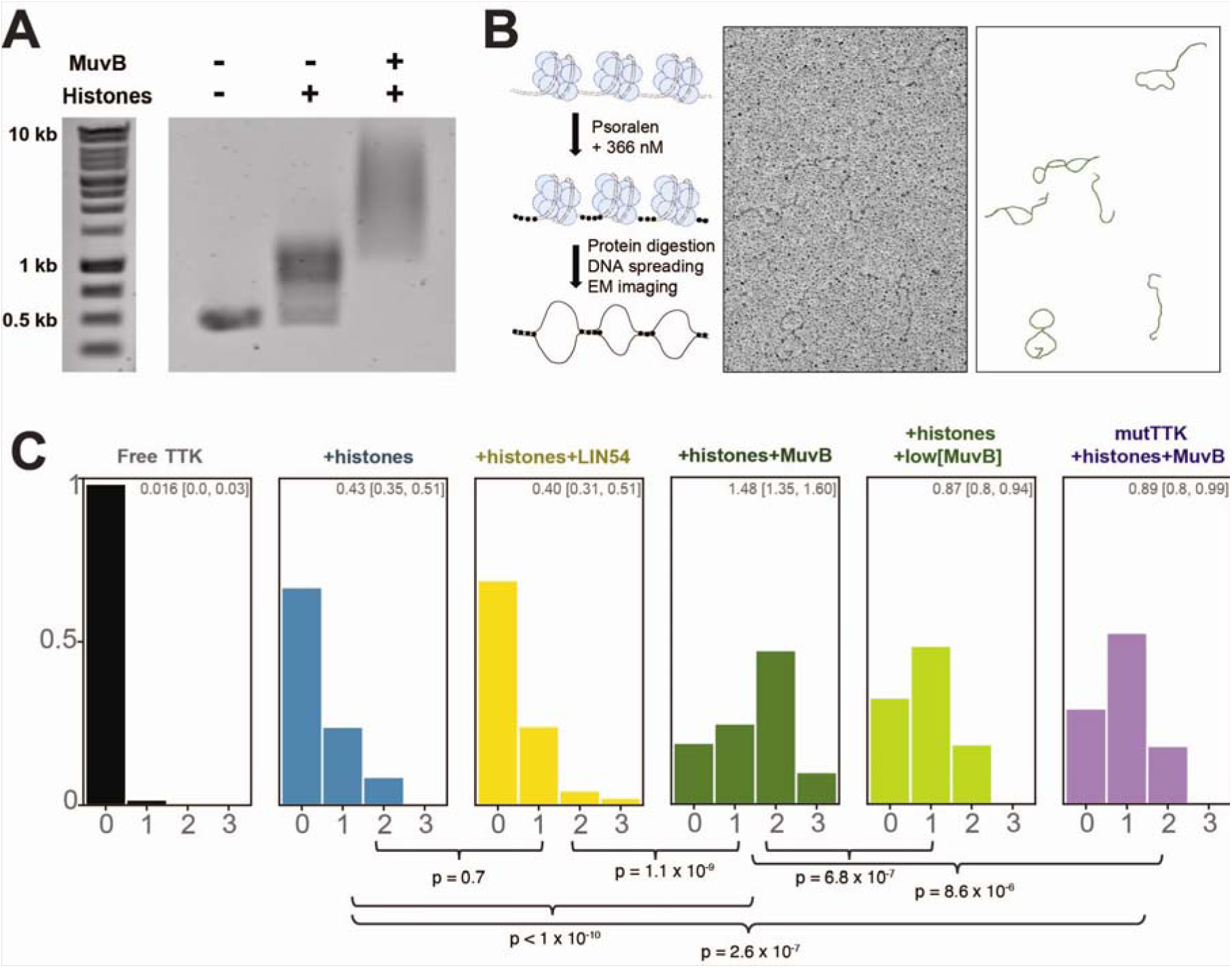
MuvB stabilizes nucleosomes on a reconstituted cell-cycle gene promoter. (A) Reconstitution of the TTK promoter with nucleosomes and MuvB. The indicated proteins were refolded with a 461 bp fragment of DNA and the samples analyzed by an agarose gel with ethidium bromide staining. (B) Schematic representation of the protocol for cross linking and imaging (left) and example electron microscopy micrograph (middle) with corresponding traced molecules (right). (C) Histograms showing the fraction of DNA molecules containing the indicated number of nucleosome-sized bubbles for a sample of 100+ analyzed DNA molecules. The MuvB concentration when used in the reaction was 1.3 μM, whereas low[MuvB] corresponds to 0.15 μM. The number at the top of the box indicates the average number of nucleosome-sized bubbles per DNA molecule with 95% confidence interval in brackets. These statistics were computed using bootstrapping with 10000 iterations of resampling. Histograms were fit to a Poisson distribution and conditions were compared using an exact-Poisson test. p-values for comparisons of average number of bubbles (λ of Poisson distribution) across conditions are reported below the histogram.

We then cross-linked our chromatinized samples and several control samples with tri-methyl psoralen, digested protein, and performed metal-shadowing electron microscopy to assess nucleosome occupancy along the DNA molecules across our conditions (Figure 5B and 5C) (Brown et al., 2015; Fei et al., 2015). In these experiments, the presence of nucleosomes is inferred from the appearance of nucleosome-sized bubbles in the micrograph (Figure S5). As expected, we observed nucleosomes in the samples folded with histone octamers prior to cross-linking but not in the sample that only contained free TTK DNA. When MuvB is present in the reconstitution reaction, we observe more molecules containing nucleosomes and an increase in the average number of nucleosomes per molecule (Figure 5C). Furthermore, the distribution of inferred nucleosomes titrated with MuvB concentration. When a lower concentration of MuvB was present, we observed fewer nucleosomes per molecule relative to the high-concentration condition. We conclude that MuvB stabilizes nucleosomes in the synthetic TTK promoter, as MuvB increased nucleosome occupancy in the equilibrium established by the reconstitution reaction. We did not observe a significant change in the nucleosome distribution with inclusion of LIN54^504-709^ alone, suggesting that binding of the CHR by the LIN54 DBD is not sufficient to increase nucleosome occupancy. When we mutated the CHR site in the TTK promoter, we observed a significant decrease in the average nucleosomes per molecule; however, this average is still greater than the average in the absence of MuvB. We propose that MuvB binds and stabilizes nucleosome occupancy in the DNA even when not bound to the CHR site through LIN54 (i.e. in *trans* association with nucleosomes) but that simultaneous engagement of both the CHR and nucleosome (i.e. in *cis* association) results in increased stability. A histone chaperone-like activity has been reported for RBAP48 in other chromatin-bound complexes, and RBAP48 binds histone octamer intermediates (Verreault et al., 1996; Zhang et al., 2013). Because our experiment probes the equilibrium established by the reconstitution reaction beginning with a folded octamer, we cannot rule out a role for MuvB in facilitating the assembly of nucleosome intermediates (Figure S5). However, we favor the interpretation that, by binding the CHR and the nucleosome (Figure 4B), MuvB stabilizes fully assembled nucleosomes in the promoter.

### MuvB associates with the +1 nucleosome in cell-cycle gene promoters

We next used an MNase ChIP approach to detect MuvB association with nucleosomes in cells (Figure 6A) (Gutin et al., 2018; Tolstorukov et al., 2013). Chromatin preparations from HCT116 cells expressing Strep-tagged LIN9 were cross-linked and MNase digested. Samples were precipitated with Strep-Tactin, and after crosslinks were reversed and protein was digested, DNA fragments were purified, ligated with bar coded adapters and sequenced. In contrast to traditional ChIP experiments that identify transcription factor binding motifs, we aimed to purify nucleosomal-DNA fragments that associate with our transcription factor (Tolstorukov et al., 2013). To enrich for these longer fragments (>100 bp), we ligated adapters after a SPRI-bead DNA purification. Compiled DNA sequences were aligned to the human genome, and we used MACS2 to locate enriched peaks corresponding to LIN9-interacting sequences.

**Figure 6:**
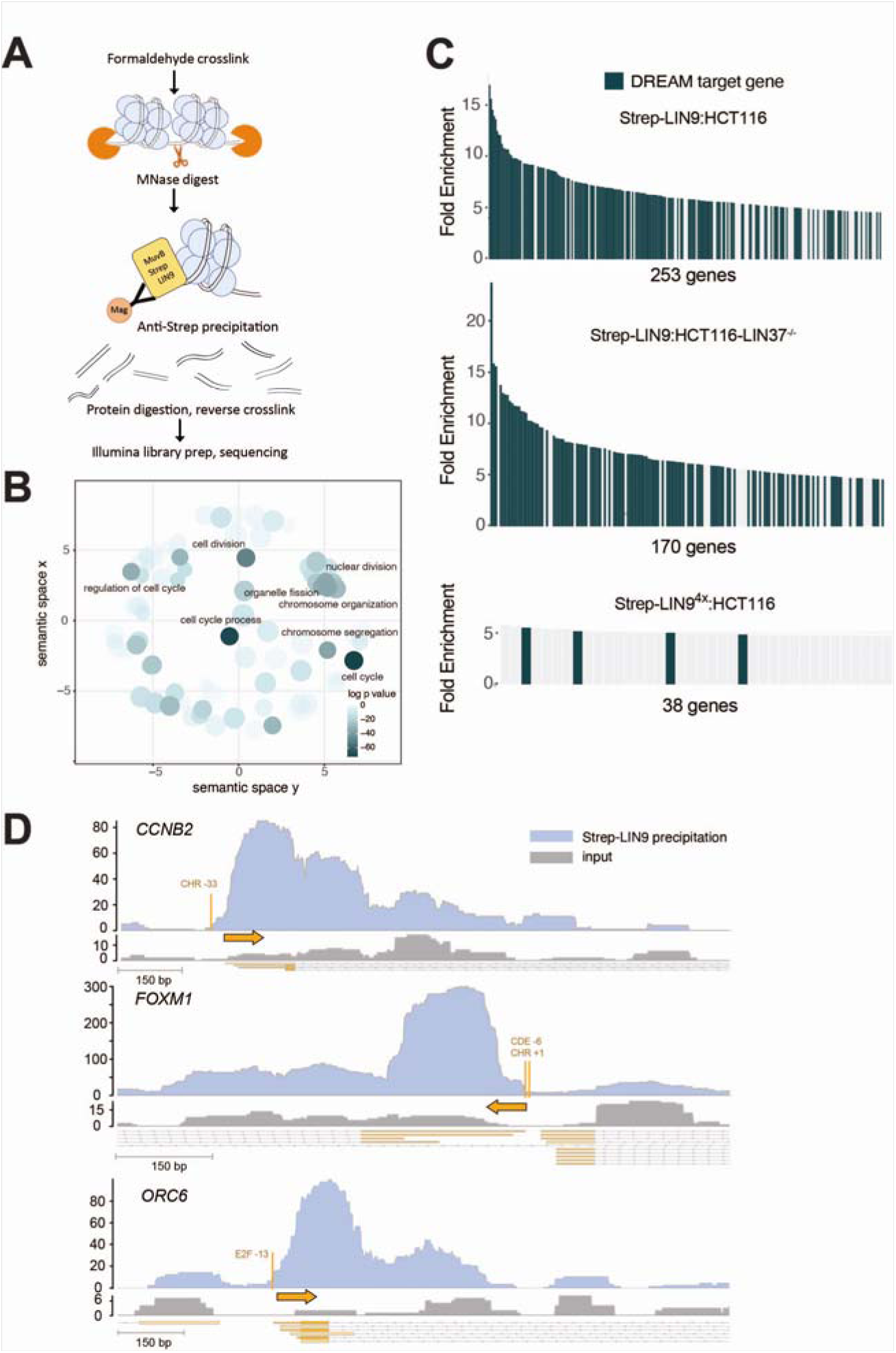
MuvB associates with nucleosomes in DREAM-regulated gene promoters in HCT116 cells. (A) Diagram of the MNase-ChIP experiment designed to enrich nucleosome-sized DNA fragments that interact with MuvB complexes. (B) Gene ontology analysis of the enriched DNA sequences following MNase digestion and precipitation from arrested cell extracts. Gene groups with p values less than 1 x 10^-30^ are labeled. (C) Top enriched genes with DNA sequences co-precipitated with Strep-LIN9 from arrested cells. The number of genes with an enrichment greater that 4.7-fold are indicated for each experiment. (D) Genome browser tracks corresponding to the *CCNB2*, *FOXM1*, and *ORC6* promoters. The number of DNA sequence reads is plotted for the input (grey) and Strep-LIN9 precipitated DNA samples. These data were for one replicate performed in HCT116 cells. Data for other replicated and experiments are shown in Figure S6. The transcription start site (TSS) in each gene (base of orange arrow) along with the position of the DREAM-binding DNA motif relative to the TSS are indicated.

We first analyzed precipitated DNA sequences from HCT116 cells that were treated with Nutlin-3a, which induces DREAM-mediated repression of both S phase and M phase genes through the p53 pathway (Figure S6A) (Schade et al., 2019a; Uxa et al., 2019). By comparing Strep-LIN9-precipitated samples to control samples in which cells were transfected with empty vector, we identified 253 genes with MACS peaks having greater than 4.7-fold enrichment in sequencing reads. Gene ontology analysis of this data set reveals enrichment in genes related to cell-cycle, mitotic division, and response to DNA damage (Figure 6B). We found that 177 (70%) of the 253 most enriched genes have previously been identified as DREAM regulated genes based on LIN9 and E2F4/p130 ChIP, RNA expression, and promoter analysis (Fischer et al., 2016; Litovchick et al., 2007; Muller et al., 2014; Uxa et al., 2019) (Figure 6C). Moreover, these DREAM genes tended to show higher enrichment than the other identified genes among the top hits. We performed two additional replicate experiments, one technical replicate with a different MNase concentration for the digestion and one biological replicate, and we found that the enrichment of many DREAM genes was reproducible (Figure S6B). We performed an analogous experiment in which we expressed Strep-LIN9^4X^. The LIN9^4x^ mutant fails to associate with LIN37 and RBAP48 but still associates with CHR consensus sites (Figures 3D, 6C, S6A, and S6C). Considering the same 4.7-fold threshold, this data set contained fewer genes overall and only four DREAM genes containing enriched sequences (Figure 6C). We conclude that our experimental protocol successfully enriches LIN9-bound DNA sequences at expected cell-cycle genes in HCT116 cells and that enrichment depends on intact MuvBN.

Inspection of the WT LIN9-immunoprecipitated sequence reads aligned to the human genome reveals enrichment of DNA corresponding to nucleosome-sized fragments (∼150 base pairs) near the transcription start site and E2F or CHR consensus sites in the DREAM-regulated genes (Figures 6D and S6D). For example, in the *CCNB2* promoter, which contains a canonical CHR DREAM-binding site, the strongest enriched peak is located just downstream of the closely spaced TSS and CHR site. This nucleosome corresponds to the +1 nucleosome, which is commonly positioned at repressed genes (Hughes and Rando, 2014; Schones et al., 2008; Struhl and Segal, 2013). We observed secondary sites of enrichment, which correspond to nucleosomes (e.g. +2 and +3 nucleosomes) further downstream of the TSS. The enrichment decreases with increasing distance from the CHR site. In *FOXM1* and *ORC6*, which contain CDE-CHR and E2F binding sites for DREAM respectively, we observed a similar pattern, with the +1 nucleosome showing the strongest enrichment, followed by a weaker coverage of the distal nucleosomes. Notably, these enriched peaks corresponding to nucleosomes are not observable in the Strep-LIN9^4x^ mutant data set (Figure S6D).

We aligned the promoter regions of the 177 enriched DREAM genes according to their transcription start sites (TSS) to identify more broadly the structural signature of LIN9-associated nucleosomes. Most of these genes show a sharply positioned nucleosome within 150 bases downstream of the TSS (Figure 7A). We conclude that LIN9 primarily immunoprecipitated the +1 nucleosome in these promoters. Considering that expressed wild-typeLIN9 forms complexes with other endogenous MuvB components (Figure 1D) but that LIN9^4X^ does not associate with LIN37 and RBAP48 (Figure 3D), we further conclude that these nucleosomes are bound by MuvB complexes and that these interactions are mediated by MuvB^N^.

**Figure 7:**
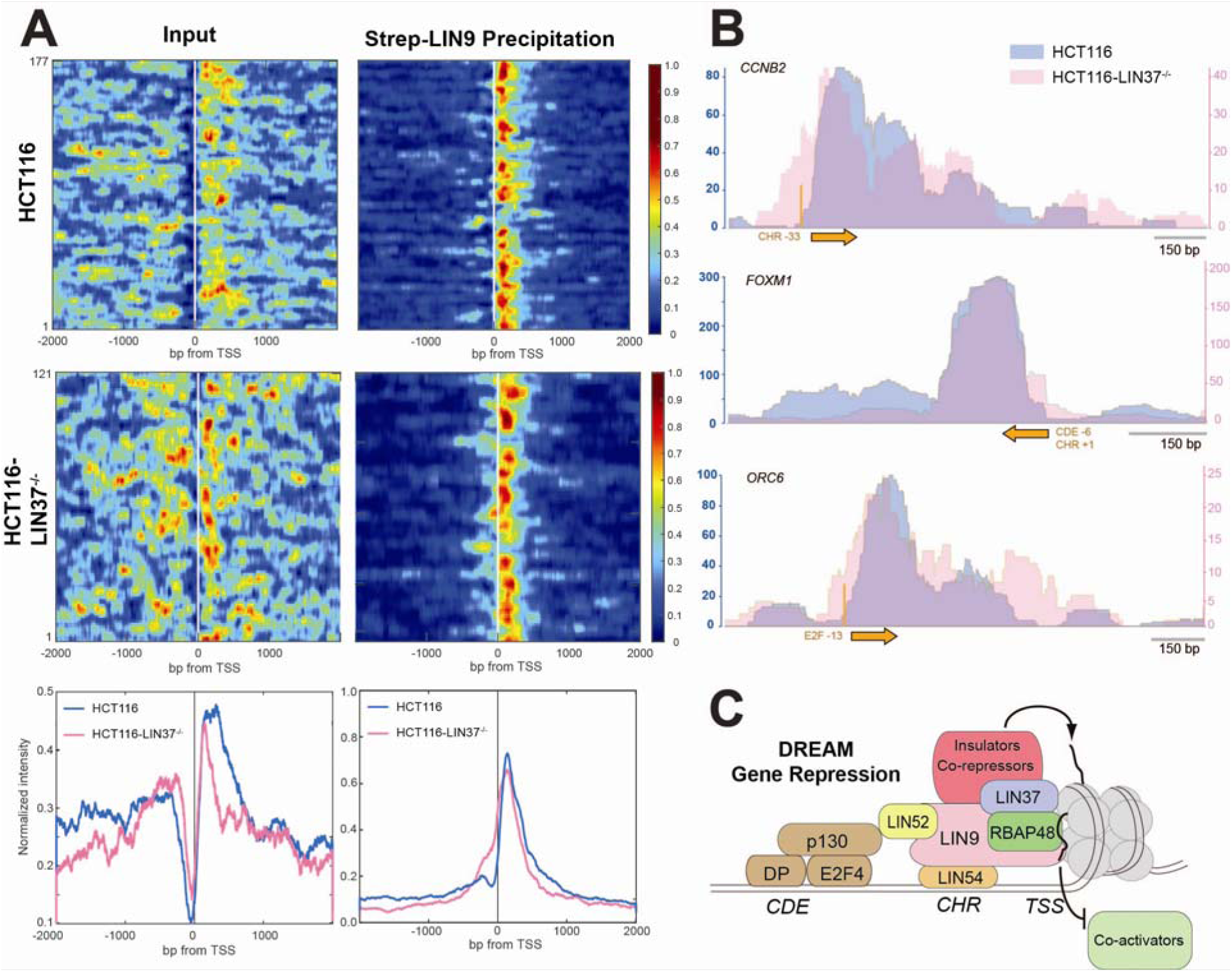
The sharp positioning of the MuvB-bound +1 nucleosome correlates with gene repression. (A) Heat maps showing relative read density across all genes that show greater than 4.7-fold enrichment in the Strep-LIN9 precipitant and correspond to a known DREAM gene. Both input and immunoprecipitant sequencing data are shown for the same gene set and in the same order. At the bottom are the aggregated and normalized read intensities across all genes shown in the heat map for the input and precipitated DNA data sets. Data from HCT116 (blue) and HCT116-LIN37^-/-^ (pink) are shown. The observation of more peaks with periodicity corresponding to nucleosomes (i.e. the +2 and +3 nucleosome peaks) in the HCT116-LIN37^-/-^ input data set is consistent with DREAM genes being active during Nutlin-3a-induced quiescence of those cells (Uxa et al., 2019). (B) Comparison of three example gene tracks showing the number of DNA sequence reads in Strep-LIN9 precipitated samples from HCT116 (blue, left axis) and HCT116-LIN37^-/-^ (pink, right axis) cells. The TSS (orange arrow) and DREAM-binding DNA motifs are indicated. (C) Overall model showing organization of DREAM and the association between MuvBN and the +1 nucleosome, which we propose mediates gene repression.

We emphasize that the enriched nucleosomes in the set of DREAM genes do not overlap with the E2F and CHR consensus binding sites, suggesting that DREAM binds these in linker DNA and not in the nucleosome core particle. Nucleosomes are positioned next to the DREAM-binding site, which is typically in close proximity to the TSS (Figure 6D), and do not necessarily contain the E2F and CHR DNA sequence motifs. We also note that the primary and secondary peaks in the sequence coverage persist when the reads are filtered for exclusively mononucleosome-sized inserts (Figure S6E). This observation that MuvB precipitated both proximal but not overlapping and distal nucleosomes to its consensus binding sequence further suggests that MuvB makes direct contact with the nucleosome core. This interpretation is consistent with our biochemical observations that interactions with nucleosomes are facilitated through protein-protein binding rather than through proximal DNA interactions (Figure 5). Importantly, the peak corresponding to the +1 nucleosome is stronger and more tightly positioned in the precipitated sequencing data compared to the input data (Figures 6D and 7A). This enrichment of a strongly positioned nucleosome is consistent with a role for MuvB in binding and stabilizing the +1 nucleosome in DREAM promoters.

### MuvB association with a tightly positioned +1 nucleosome correlates with gene repression

We next performed a similar MNase-ChIP experiment using LIN37 knock-out HCT116 cells (HCT116-LIN37^-/-^, Figures 6C, 7A, 7B, S6A, and S6D). In these cells, the MuvB complex assembles on CHR promoters, but cell-cycle genes are no longer fully repressed by DREAM when cells are arrested with Nutlin-3a (Mages et al., 2017; Uxa et al., 2019). We in fact observed in the input MNase data from the set of DREAM genes the nucleosome phasing pattern that is characteristic of genes undergoing transcription (Figure 7B) (Oruba et al., 2020; Schones et al., 2008). We still observed enrichment of known DREAM genes in the pool of Strep-LIN9 precipitated DNA reads (Figure 6C), and many of the enriched genes overlap between the data sets from wild-type and LIN37 knock-out cells (Figure S6B). We note that this result from precipitating Strep-LIN9 from knock-out cells is distinct from what we observed precipitating Strep-LIN9^4x^ from wild-type cells. In the former experiment LIN37 is missing from MuvB complexes, while in the latter both LIN37 and RBAP48 are missing; however, both complexes can associate with DNA (Figures 3D and S6C). From this comparison, we conclude that RBAP48 is necessary for nucleosome association.

Analysis of nucleosome occupancy generated for enriched genes in the data set from knock-out cells suggests that MuvB still associates with +1 nucleosomes (Figures 7A, 7B, and S7). However, the bound nucleosomes are distributed over a broader region of DNA, i.e. the boundaries of the positioned nucleosome are more poorly defined, and the position more typically encroaches on the TSS. While we cannot determine that this broader distribution of nucleosome positions is directly a result of the absence of LIN37 from the complex or is a signature of expressed DREAM genes in the KO cells, these results demonstrate that the sharply positioned MuvB-associated nucleosomes in the WT cells correlate with gene repression.

## Discussion

Genetic studies across model organisms all point to the function of the MuvB core as an intrinsically repressive complex that interacts with other TFs to modulate gene expression. In *C.elegans*, even when the p107/p130 ortholog is knocked-out such that DREAM does not form on promoters, the MuvB core retains the ability to repress target genes (Goetsch et al., 2017). In *Drosophila,* the lethal *myb-null* phenotype can be rescued by the loss of function of the fly orthologs of LIN9 and LIN37, which restores expression of Myb target genes (Beall et al., 2004; Beall et al., 2007; Wen et al., 2008). In mammalian cells, LIN37 knock-out and LIN9 knock-down leads to specific loss of repression of cell-cycle genes upon driving cell-cycle exit (Litovchick et al., 2007; Mages et al., 2017; Reichert et al., 2010), and a similar defect is observed upon loss of the RBAP48 ortholog in *Drosophila* (Taylor-Harding et al., 2004). While mammalian LIN9 loss also results in failure to activate mitotic genes, this activation defect may be linked to the requirement of LIN9 for recruitment of B-Myb (Guiley et al., 2018; Litovchick et al., 2007; Osterloh et al., 2007; Reichert et al., 2010). Together, these results demonstrate that LIN9, LIN37 and RBAP48 contribute to a repressive MuvB function.

Our results implicate the MuvBN subcomplex as the structural unit in MuvB responsible for this intrinsic repressive function and link repression to nucleosome binding. The structure and biochemical data demonstrate that LIN9 and LIN37 form a scaffold for the MuvB core in that they bind and assemble LIN52, LIN54 and RBAP48 (Figure 7C). The MuvBN structure contains RBAP48 and the conserved sequences of LIN37 that are both required specifically for cell-cycle gene repression. Our data demonstrate that MuvB binds and stabilizes nucleosomes and that MuvBN, which contains the repressor subunits, is sufficient for this interaction. We observed association of LIN9-containing MuvB complexes with the +1 nucleosome in the promoters of repressed cell-cycle genes in arrested cells, but this association is lost with a LIN9 mutant that does not assemble the MuvBN components LIN37 and RBAP48. While we still see association of MuvB with nucleosomes in active gene promoters in arrested LIN37 knock-out cells, the associated nucleosome is more strongly positioned under conditions of repression. We propose that MuvB association with +1 nucleosomes contributes to repression by inhibition of remodeling, polymerase activity, or posttranslational histone modification required for transcription (Figure 7C). For example, association of MuvB with the histone H3 tail may sequester the tail from other chromatin binding proteins and remodelers. Another possibility is that by tightly binding and positioning the +1 nucleosome, MuvB might increases the energy barrier that stalls RNA polymerase activity, resulting in uninitiated or aborted transcripts. By binding through multiple modes, i.e. histone tails and core, MuvB could prevent the unwrapping and movement of the +1 nucleosome. Although initial mass spectrometry analysis revealed few binding partners to human MuvB that could explain its repressive role, more recent studies have found MuvB can in certain contexts recruit proteins such as Sin3A (Bainor et al., 2018; Kim et al., 2021). Additional factors may also function to enhance repression in addition to the nucleosome binding activity of MuvB. Our result that MuvB can bind nucleosomes even in the absence of an H3 tail interaction suggests that the H3 site in RBAP48, which our structure shows is accessible in MuvB, might be used by MuvB to recruit other repressor complexes.

Research on the structure of chromatin has revealed important factors that determine the nucleosome position in the genome, including intrinsic properties of DNA sequence, chromatin remodeling complexes, the polymerase machinery, and sequence specific TFs (Kujirai and Kurumizaka, 2020; Lai and Pugh, 2017; Teves et al., 2014). The role of TFs has focused on their potential for maintaining the nucleosome depleted region around the TSS and for establishing the +1 nucleosome. Evidence supports a model in which the mutually exclusive interaction between TFs and histones for DNA allows TFs to act as a barrier for nucleosome deposition such that the +1 and other proximal nucleosomes form at the closest accessible sites. Our data support a more direct function for TFs in establishing the +1 position through physical association and correlate this association in cells with a more tightly positioned nucleosome at repressed genes. Under conditions when MuvB is not actively repressing (LIN37 KO cells in quiescence), we observe more variability in the nucleosome position. The extent to which these observations result from the dynamics of RNA polymerase during the transition from repressed to active genes remains uncertain.

Several important questions remain about this MuvB repressive function including the structural mechanism of nucleosome recognition and the role of LIN37. RBAP48 in many studies is sufficient for nucleosome binding, and it is still present in MuvB complexes that lack LIN37 yet cannot repress gene expression (Mages et al., 2017). We speculate that this non-functional complex may be unstable or improperly structured such that it cannot enact repression or bind co-repressors. The extensive interaction interface and co-dependence of their association in our mutagenesis study support the hypothesis that the core subunits of MuvBN, (LIN9, LIN37 and RBAP48) co-fold to form a stable complex. Another important remaining question is how the structure and function of MuvB changes such that it switches from a repressor to an activator of gene expression once cells enter the cell-cycle. MuvB components are still present on the promoter and are required for recruiting B-Myb and FoxM1. One possibility is that MuvB repression activity is relieved, for example by binding of B-Myb, which is consistent with observations in *Drosophila* that the MuvB-binding sequence in B-Myb is alone sufficient to rescue a B-Myb deletion phenotype (Andrejka et al., 2011). Another possibility is that Cdk phosphorylation, detected on all the MuvB subunits, plays a role in modulating MuvB function (Odajima et al., 2016). A third possibility is that in addition to its repressive function, MuvB positions the +1 nucleosome to prime genes for expression upon the binding of the activator transcription factors B-Myb and FoxM1. In this mechanism, MuvB may facilitate the acetylation of histones by the p300 acetylation machinery, which is recruited by the activator TFs. Through TF and p300 association, MuvB may also help recruit the basal transcription machinery. Finally, it will be important to understand how widespread interactions of TFs with the +1 nucleosome are and how these interactions regulate chromatin and gene expression

## Methods

### Plasmids for protein expression

The LIN9 and RBAP48 ORFs were amplified from cDNA derived from mouse NIH3T3 cells by standard PCR. The EGFP ORF was amplified from pEGFP-N1 (Clontech). The ORFs were cloned into pcDNA3.1(+) and fused either with an N-terminal 3xFlag tag (Lin9) or and N-terminal 1xFlag tag (RBAP48, EGFP). Site directed mutagenesis was performed following the QuikChange protocol (Stratagene). For MNase-Seq experiments, the Lin9 ORFs were subcloned into pcDNA3.1(+) containing and N-terminal Twin-Strep tag.

### Protein expression

To assemble the entire MuvB complex, proteins (GST- or Strep-LIN9^94-538^, GST-LIN37, GST-LIN52, His- or GST-LIN54^504-749^, and Strep-RBAP48) were co-expressed in Sf9 cells via baculovirus infection. Cell pellets were harvested after 72 hours growth in suspension at 27°C, and complexes were purified using GST-affinity purification followed by Strep-affinity purification. After removal of affinity tags through TEV protease cleavage, purified complexes were isolated through size-exclusion chromatography using a Superdex200 column. The final buffer contained 200 mM NaCl, 25mM Tris HCl and 1mM DTT at pH 8.0. The MuvBN subcomplex was assembled by co-expressing GST-LIN9^94-274^, LIN37^92-130^, and full length RBAP48 in Sf9 cells as described for the full complex. The subcomplex was purified using GST affinity purification followed by anion exchange. Affinity tags were then removed with TEV protease and complex was isolated with a Superdex 200 column. The final buffer contained 150 mM NaCl, 25mM Tris HCl and 1mM DTT at pH 8.0.

### X-ray crystallography

The MuvBN subcomplex was crystallized in a sitting drop at 4°C containing 0.2 M sodium tartrate tetrahydrate, 0.1 M bis-tris propane pH 6.5, and 20% PEG 3350. Crystals were harvested and directly frozen in liquid nitrogen. Data were collected at λ = 1.0332Å and 100 K on Beamline 23-ID-B at the Advanced Photon Source, Argonne National Laboratory. Diffraction spots were integrated with Mosflm (Leslie, 2006) and scaled with SCALE-IT (Howell and Smith, 1992). Phases were solved by molecular replacement with PHASER (Mccoy et al., 2007) and using RBAP48 (PDB: 3GFC) was as a search model. The initial model was rebuilt with Coot (Emsley and Cowtan, 2004), and LIN9 and LIN37 were added to the unmodeled electron density. The resulting model was refined with Phenix (Adams et al., 2010). Several rounds of position refinement with simulated annealing and individual temperature-factor refinement with default restraints were applied. The final refined model was deposited in the Protein Data Bank under Accession Code PDB ID: 7N40.

### Co-immunoprecipitation and DNA affinity experiments

Human HCT116 colon carcinoma cells were cultivated in DMEM supplemented with 10% FBS. Transfections were performed in 10cm plates using 7 ug plasmid and 35 ul PEI per plate. To stimulate DREAM formation, cells were treated with 10uM Nutlin-3a for 24 hours. Cells were harvested 48h after transfection. Whole cell extracts were prepared by lysing the cells in IP lysis buffer (50 mM TRIS-HCl pH 8.0, 0.5% Triton-X 100, 0.5 mM EDTA, 150 mM NaCl, 1mM DTT, and protease inhibitors) for 10 min on ice followed by 5x 1s direct sonication. Flag tagged proteins were immunoprecipitated from 2-3mg cellular extracts with Pierce Anti-DYKDDDDK Magnetic Agarose (Invitrogen). Beads were washed 5x with 1ml IP lysis buffer end eluted with 50µl 1xLaemmli buffer. 12µg of input samples and 12µl IP samples were analyzed by SDS-PAGE and western blot following standard protocols. The following antibodies were applied for protein detection: FLAG-HRP (RRID:AB_2017593, Santa Cruz Biotechnology), p130/RBL2 (D9T7M) (RRID:AB_2798274, Cell Signaling), LIN54 A303-799A (RRID:AB_11218173, Bethyl Laboratories), LIN9 ab62329 (RRID:AB_1269309, Abcam), RBBP4 A301-206A (RRID:AB_890631, Bethyl Laboratories), LIN37-T3 (custom-made at Pineda Antikörper-Service, Berlin, Germany) (Muller et al., 2017).

For DNA affinity experiments,

### Nucleosome reconstitution

Xenopus histones as well as their tailless counterparts were expressed and purified in E.coli as inclusion body preparations as previously (Luger et al., 1999; Yang and Narlikar, 2007). Octamer reconstitution was completed by mixing equimolar amounts of purified histones in a buffer containing 7 M guanidinium HCl, 20 mM Tris pH 7.5 and 10mM β followed by dialysis into 2 M NaCl, 10 mM Tris pH7.5, 1 mM EDTA and 5 mM β mercaptoethanol. Folded octamers were purified using size-exclusion chromatography on a Superdex 200 column. Nucleosome reconstitution was performed by mixing purified histone octamers with the Widom 601 positioning sequence and de-salting by gradient dialysis (Luger et al., 1999). For Widom nucleosomes, we used a 1.1:1 ratio of octamers:DNA molecules. At a salt concentration of 50 mM, nucleosome samples were collected in a buffer containing 50 mM Tris pH 7.5 and 1 mM DTT.

### Florescence polarization assay

Histone peptides were synthesized with fluorescein. For experiments with Widom nucleosomes, the 601 sequences were PCR amplified with a primer containing fluorescein and reconstituted with octamer as described above. 20 nM peptide was mixed with varying concentrations of MuvB protein complex in a buffer containing 50 mM Tris pH 7.5, 150 mM NaCl, 1 mM DTT, and 0.1% (v/v) Tween-20. Twenty microliters of the reaction were used for the measurement in a 384-well plate. Fluorescence polarization (FP) measurements were made in triplicate, using a Perkin-Elmer EnVision plate reader. The K_D_ values were calculated using Prism 8 (Version 8.2.1).

### Electron microscopy on reconstituted promoters

The minimal region of the human *TTK* promoter (461bp) was cloned, amplified by PCR, and purified by agarose gel extraction. Histone octamers were folded with the *TTK* DNA as described above for Widom nucleosomes, but we used an octamer to DNA ratio of 3.1:1 to allow for the formation of di- and tri-nucleosome species. For the relevant conditions, purified MuvB complex or LIN54 was added to the nucleosome folding reaction during the de-salting process at a NaCl concentration of ∼800mM. Cross-linking of gene promoters and electron micrograph preparation was performed using the protocol outlined in (Brown et al., 2015). In brief, samples were treated with trimethylpsoralen and UV radiation to allow double-stranded DNA crosslinks to form at unprotected, octamer free regions. Following crosslinking, proteins were digested by Proteinase K, and DNA molecules were purified, denatured, and spread across the surface of a copper transmission electron microscopy grid. Electron micrographs of all samples were prepared by rotary metal shadowing, and grids were visualized and collected on a JEOL 1230 TEM at the UC Santa Cruz IBSC Microscopy facility and a Tecnai 12 TEM at the UC Berkeley ELM lab. DNA molecules were traced, and molecules coordinates were saved using Fiji tools as previously described (Brown et al., 2015). Resulting traces were analyzed using custom python tools. Each DNA strand was traced such that an “end” of the molecule could be identified. Thus, every coordinate in one strand can be aligned to its complement by closest distance. Coordinates are assigned to base positions using a scale based on the physical distance between coordinates within each strand based on the known length of the TTK promoter (461 bp). A base pair is labeled single stranded if the distance between strands exceeds a threshold distance, determined empirically. Once all base pairs are labeled, “bubbles” are determined by contiguous single stranded stretches. Finally, a single stranded “bubble” is labeled a nucleosome if its length is >90 basepairs (Figure S5). Bubble fusions occur such that two or more adjacent nucleosomes form one contiguous bubble; for this analysis, bubble fusions were labeled as a single nucleosome. This estimate was used because the number of total bubble fusions observed within the dataset was small <5%.

### MNase-ChIP

HCT116 cells transfected with Strep-Lin9 constructs or an empty Strep expression plasmid. 24h after transfection, cells were treated with 10µM Nutlin-3a (Selleckchem) and harvested after 48h. Cells were crosslinked with 1% formaldehyde for 15 minutes. The crosslinking reaction was stopped with glycine, cells were washed twice with PBS, and pellets were collected. Cell lysis and MNase digestion (1x or 5x) were performed as described earlier, and following digestion, LIN9-bound samples were precipitated using Streptactin-XT magnetic beads (IBA Lifesciences) (Gutin et al., 2018). Both input and IP samples were subject to RNAse treatment and proteinase K digestion and were reverse crosslinked by incubation at 65 °C for 16 hours. DNA was purified by 2x SPRI bead clean-up. Library prep was carried out using NEB Next Ultra II kits, and paired end sequencing was carried out on the NovaSeq 150bp platform for Illumina at Novogene Biotech, Co., LTD.

Sequencing reads were aligned against hg38 using the bwa-mem aligner (Karolchik et al., 2004; Quinlan and Hall, 2010). Samtools and bedtools were using to convert data into bam and bed files respectively. Peak calling for the precipitated samples was performed using the MACS2 - *bampe* algorithm and using the empty Strep-IP conditions as the control. To retrieve gene names for MACS2 peaks, coordinates were intersected with known genes using the Table Browser tool provided by UCSC genome browser. Gene ontology analysis was performed on MACS2 peaks showing a greater than 4.7-fold enrichment using the web-based tools GeneOntology.org and Revigo (Supek et al., 2011). We generated coverage plots of our reads using Gviz and rtracklayer and other opensource R tools.

We utilized NucTools in paired-end mode to analyze nucleosome occupancy on input and Strep-LIN9 precipitated reads with single base-pair resolution (bin width=1bp) (Vainshtein et al., 2017). We restricted our analysis to genes that showed a MACS enrichment of greater than 4.7-fold and were previously annotated to bind DREAM, to respond to p53 stimulation, and to become derepressed in LIN37 knockout cells (Fischer et al., 2016; Litovchick et al., 2007; Muller et al., 2014; Uxa et al., 2019). We retrieved the TSS for this set of genes, either from those annotations or using bioMart (Smedley et al., 2009). As needed, the TSS sites were mapped on to hg38 using liftover. We oriented the output to center on the TSS and maintain a uniform direction of transcription. We then utilized the Cluster Map Builder feature of NucTools to generate aggregate plots and heatmaps of our genes.

## Acknowledgements

This work was supported by a grant from the National Institutes of Health to S.M.R. (R01GM124148). GM/CA@APS has been funded by the National Cancer Institute (ACB-12002) and the National Institute of General Medical Sciences (AGM-12006, P30GM138396). This research used resources of the Advanced Photon Source, a U.S. Department of Energy (DOE) Office of Science User Facility operated for the DOE Office of Science by Argonne National Laboratory under Contract No. DE-AC02-06CH11357. The Eiger 16M detector at GM/CA-XSD was funded by NIH grant S10 OD012289. We thank the staff at the University of California Berkeley Electron Microscope Laboratory for advice and assistance in electron microscopy sample preparation and data collection. We thank Joseph Lipsick and Geeta Narlikar for histone plasmids.

## Supplementary Figures

**Table S1:** Data collection and Refinement statistics. Data were collected on a single crystal. Values in parentheses are for the highest resolution shell.

| Data collection |  |
| --- | --- |
| Beam line | APS(23IDB) |
| Resolution Range (Å) | 60.0-2.55 (2.69 - 2.55) |
| Space group | C 1 2 1 |
| Unit cell dimensions(a,b,c) (Å), ($\alpha$, $\beta$, $\gamma$) (°) | 133.58 77.81 64.56<br>90 114.71 90 |
| Wavelength (Å) | 1.03 |
| Total observations | 43599 (6521) |
| Unique reflections | 19131 (2805) |
| Completeness (%) | 97.4 (97.8) |
| $R_{\text{merge}}$ | 13.5 (52.1) |
| $\langle I/\sigma \rangle$ | 7.9 (3.5) |
| $CC_{1/2}$ | 0.97 (0.47) |
| Redundancy (highest shell) | 2.3 (2.3) |
| Refinement |  |
| $R_{\text{work}}$ %/ $R_{\text{free}}$ % | 16.1/ 25.0 |
| Number of non-hydrogen atoms | 4862 |
| Protein | 4720 |
| Water | 142 |
| Wilson B-factor (Å<sup>2</sup>) | 32.79 |
| RMSD Bond length (Å) | 0.008 |
| RMSD Bond angle (°) | 0.94 |
| Ramachandran favored(%) / Ramachandran outliers (%) | 96.0/0.0 |

**Figure S1:**
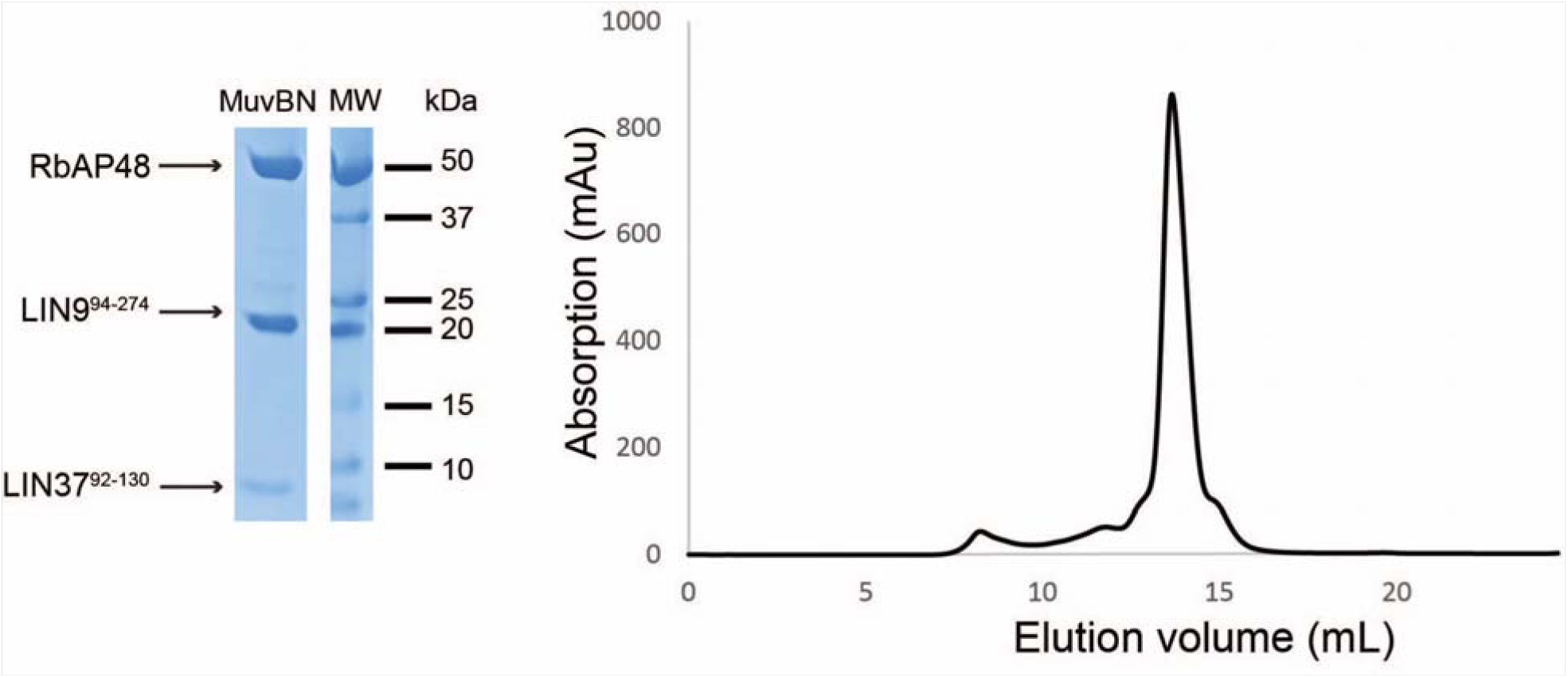
MuvBN reconstitution for crystallization. Stable complex of RBAP48 and the indicated LIN9 and LIN37 constructs eluted on Superdex 200. (Left) Coomassie stained gel of three subunits following purification. (Right) UV absorption trace from size-exclusion chromatography.

**Figure S2:**
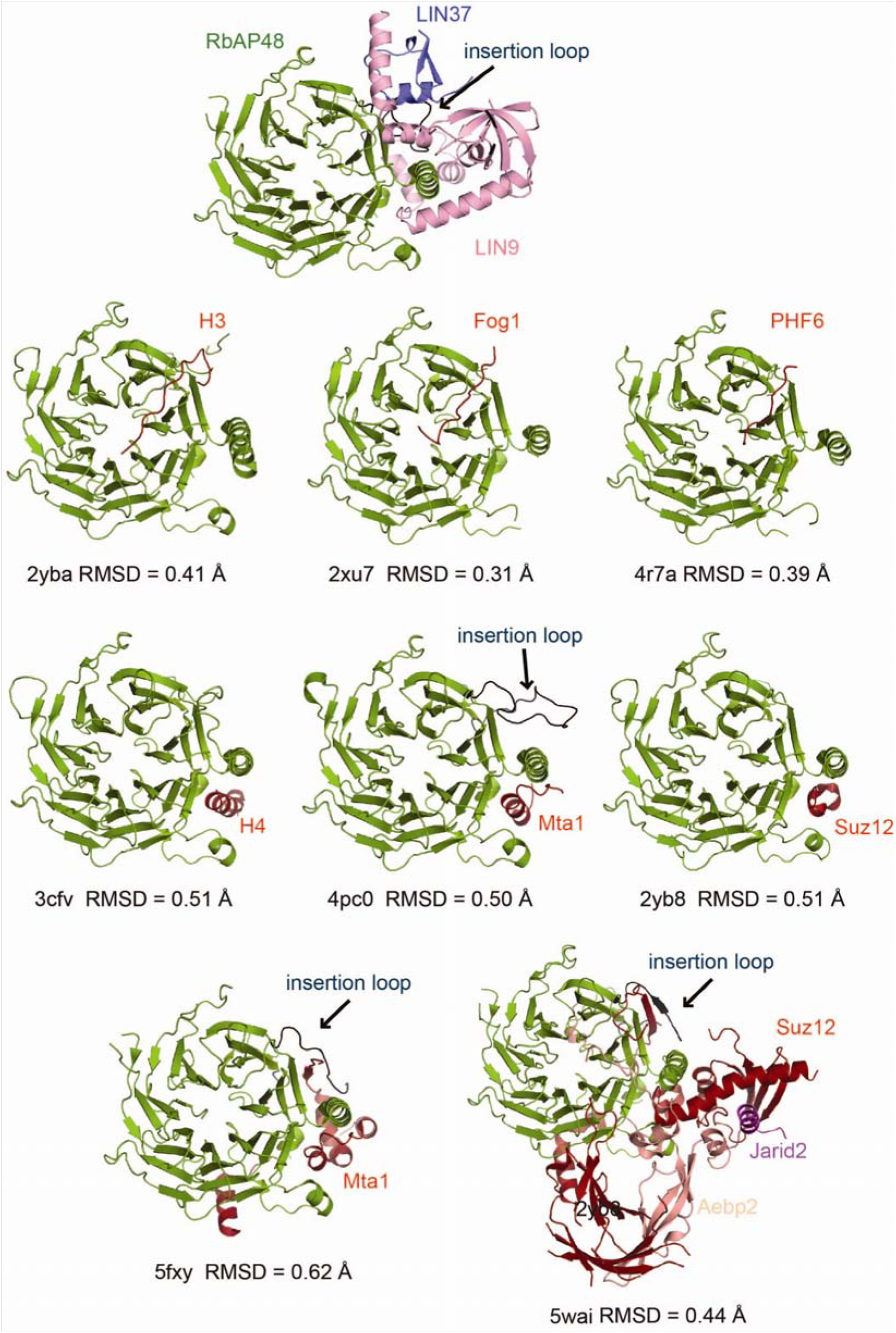
Comparison of RBAP48 structures and interactions with different peptides and proteins. The MuvBN complex structure is in shown at the top. Pairwise RMSD for C_α_ atoms in RBAP48 are indicated under each structure. Structures with peptides (top two rows) show two major binding sites. The H3 binding site across the face of the β-propeller domain (top row) is accessible in MuvBN, while the H4 binding site (middle row) is not. The RBAP48 insertion loop (residues 88-115) is not ordered and modeled in nearly all the structures of RBAP48 with small peptides. It appears in the structure with the Mta1 peptide, although in that crystal it is involved in packing. In the structures of larger complexes (bottom row), part of the insertion loop is ordered and forms a strand and packs against an added strand from an RBAP48-interacting partner. In the MuvBN structure, the loop is completely ordered and makes extensive interactions with both LIN9 and LIN37.

**Figure S3:**
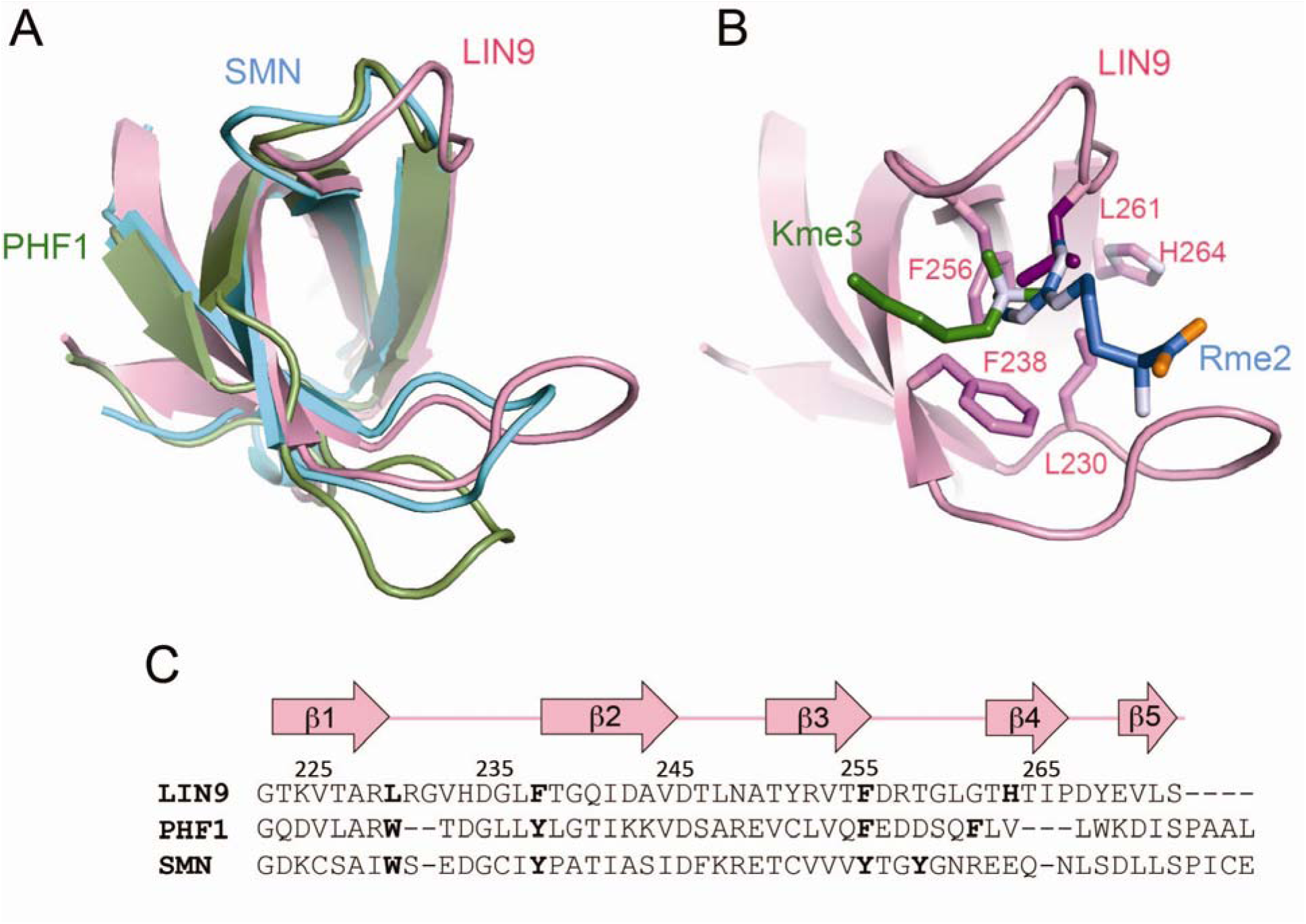
Structure of the LIN9 Tudor domain and comparison to other Tudor domains. (A) Alignment of LIN9 Tudor domain with PHF1, which binds a trimethylated peptide in histone 3 (H3K36Me3), and SMN, which recognizes a dimethylated arginine. Pairwise RMSD for C_α_ atoms are 1.0 Å between LIN9 and PHF1 and 0.9 Å between LIN9 and SMN. (B) View of LIN9 residues L230, F238, F256 and H264, which correspond to residues in the other structures that form the aromatic cage. The key modified interacting residue with the PHF1 and SMN structures are modeled in the view. The “cage” in LIN9 contains fewer aromatic residues (alignment shown in (C)), and the position of the interacting residue is occupied by L261 in LIN9. We conclude from the structure that the Tudor domain of LIN9 is non-canonical, and it may not interact with modified histone species, or it may interact in a different manner as previously observed.

**Figure S4:**
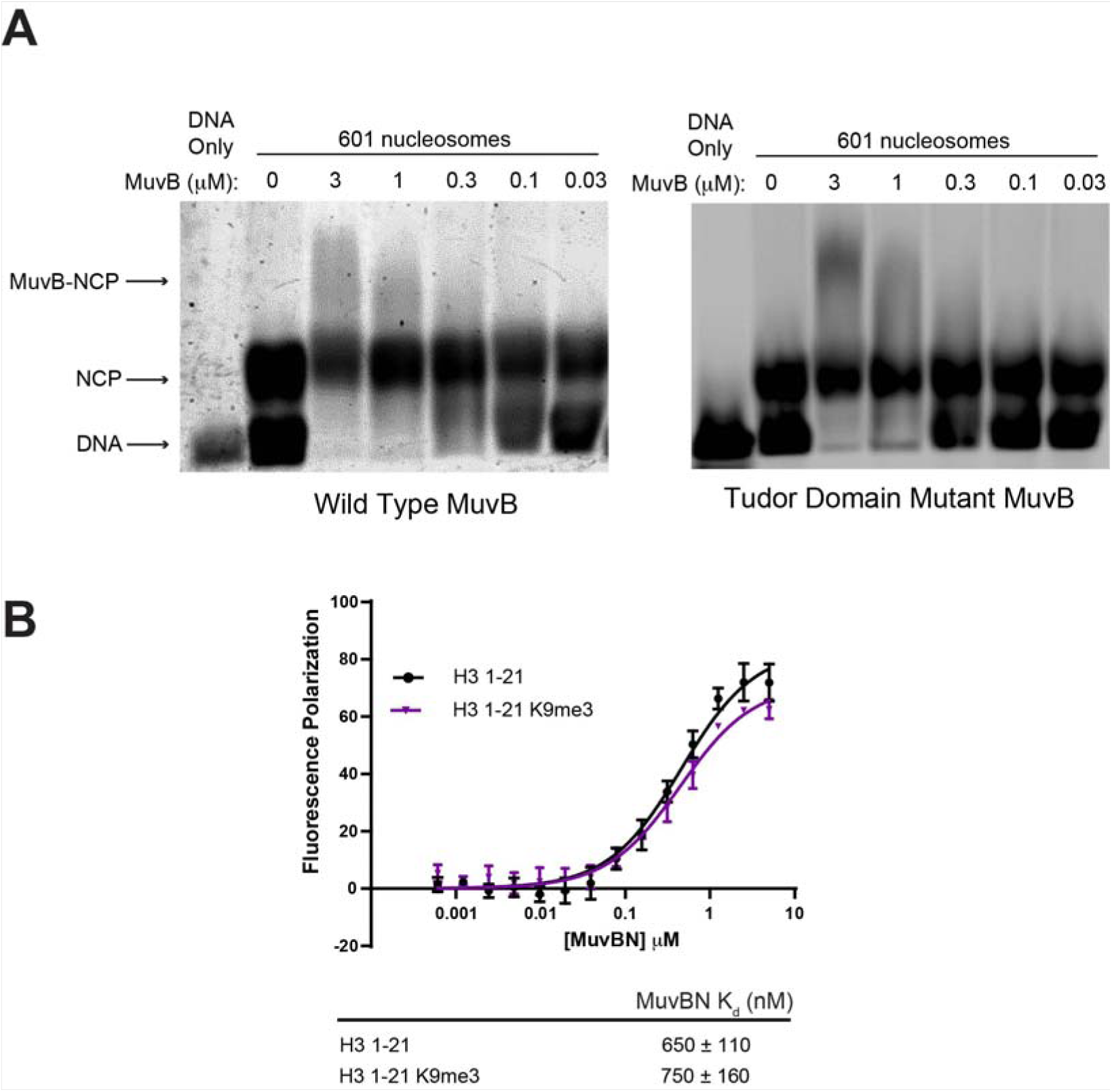
Supporting data characterizing MuvB association with nucleosomes. (A) Association of MuvB with Widom 601 nucleosomes in an electromobility shift assay. Nucleosome core particles (NCPs) were reconstituted using recombinant histone octamers and fluorescein-labeled 601 DNA. DNA alone, NCPs, and NCPs incubated with increasing concentration of MuvB was loaded onto a 0.5% agarose gel in buffer containing TBE. We reproducibly observe an increase in the ratio of NCP to free DNA when MuvB is added at concentrations near the K_d_ determined in our fluorescence polarization experiment (Figure 4). We suspect that the disappearance of the free DNA band is from a stabilizing effect of MuvB on the nucleosome. This interpretation is consistent with the lack of binding free DNA and the stabilizing effect of MuvB also observed in the data in Figure 5. An additional shift in the NCP band is also observable, particularly at higher MuvB concentrations. Wild-type MuvB and MuvB with mutations to the aromatic cage residues in the LIN9 Tudor domain show similar binding to the NCP in this assay. The LIN9 Tudor aromatic cage mutations are L230A, F238A, F256A, and H264A (Figure S3). (B) Fluorescence polarization (FP) assay as in Figure 4 but using the MuvBN subcomplex. FP is normalized to dye-labeled probe alone. Average affinities from three replicates are shown.

**Figure S5:**
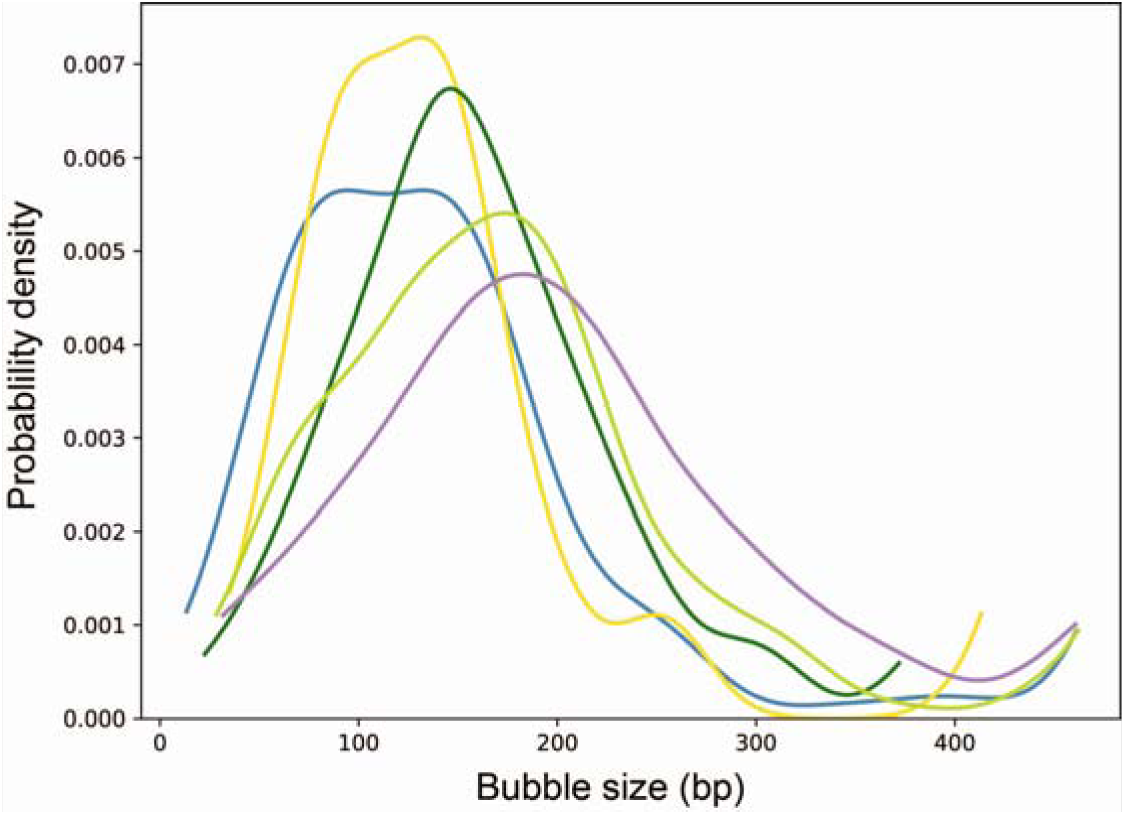
Supporting data for nucleosome crosslinking experiment with reconstituted *TTK* promoter. Estimated probability density functions for bubble size for each of the experimental conditions: blue (*TTK* promoter with histones), gold (+LIN54), dark green (+MuvB), light green (+ low[MuvB]), and purple (mutant *TTK* with histones and MuvB). Considering the resolution of the experiment and allowing for the possibility of partially wrapped octamers, we interpreted bubbles larger than 90 bp as nucleosomes. We have previously shown that a partially unwrapped pre-nucleosomal structure can be distinguished from a fully wrapped nucleosome with this psoralen cross-linking assay (Fei et al., 2015). We note that probability density around 90 bp decreases when MuvB but not LIN54 is added to the sample. This shift in the probability density away from pre-nucleosomal to nucleosomal size suggests that MuvB stabilizes the fully wrapped nucleosomes.

**Figure S6:**
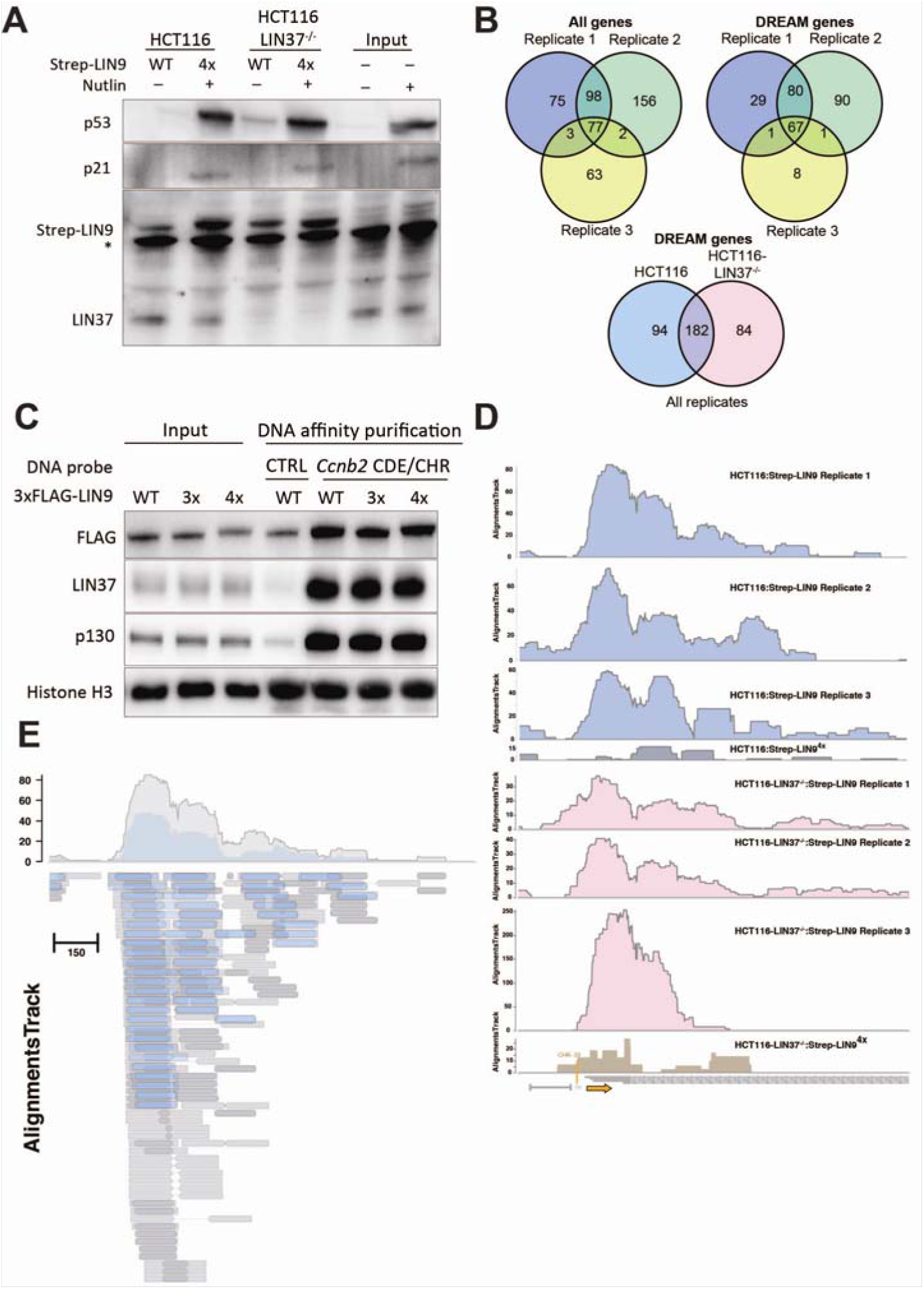
Data supporting the MNase-ChIP experiment. (A) Western blot demonstrates the expression of Strep-LIN9 and Strep-LIN9^4x^, the absence of LIN37 in the HCT116-LIN37^-/-^ cells, and the induction of p53 and the p53 target p21 upon Nutlin-3a treatment. * Indicates a nonspecific band in the blot. (B) Overlap of enriched genes between replicates of the MNase-ChIP experiment. The top two Venn diagrams show overlap of all genes (left) and DREAM genes (right) that were enriched >4.7-fold in the three replicate experiments in which Strep-LIN9 was expressed in HCT116 cells. The bottom Venn diagram shows overlap of genes that were enriched in experiments expressing Strep-LIN9 in HCT116 and HCT116-LIN37^-/-^ cells. This comparison uses the total DREAM gene list compiled across all three replicates for each condition. (C) LIN9 mutants that do not assemble LIN37 and/or RBAP48 still bind DNA containing the consensus CDE-CHR site. HCT116 cells were transfected with plasmids expressing 3xFLAG-tagged wild-type LIN9, LIN9^3x^, and LIN9^4x^. Cells were arrested with Nutlin-3a and MuvB complex components were purified. Purification was performed with a fragment of the pGL4.10 vector containing the mouse Ccnb2 CDE/CHR MuvB-binding site. Protein binding to a pGL4.10 fragment without this element was analyzed to control for background binding (CTRL). Binding of Flag-LIN9 and endogenous LIN37 and p130 was tested by western blotting. Histone H3 was probed as a control for DNA affinity purification. (D) Number of DNA sequence reads across the *CCNB2* gene track for all eight indicated experiments. The plots show DNA reads from the Strep-precipitated samples. The TSS (orange arrow) and position of the CHR element are shown. The number of reads from the Strep-LIN9^4x^ experiments are much lower and do not consistently show peaks corresponding to the +1 nucleosome. (E) DNA sequence reads from the Strep-LIN9:HCT116 precipitation are plotted for the *CCNB2* gene track as in Figure S6D. The alignment tracks of the sequence fragments are diagrammed below. In these tracks, explicitly sequenced DNA is shown as a block and inferred sequence from the paired end analysis is shown as a line. The grey tracks and coverage plot represent the full data as shown in Figure 6D and Figure S6D. The blue tracks and coverage plot represent a subset of that data, in which the alignment tracks are filtered to include only sequence reads that are 130-200 base pairs, corresponding to mononucleosome-size fragments.

**Figure S7:**
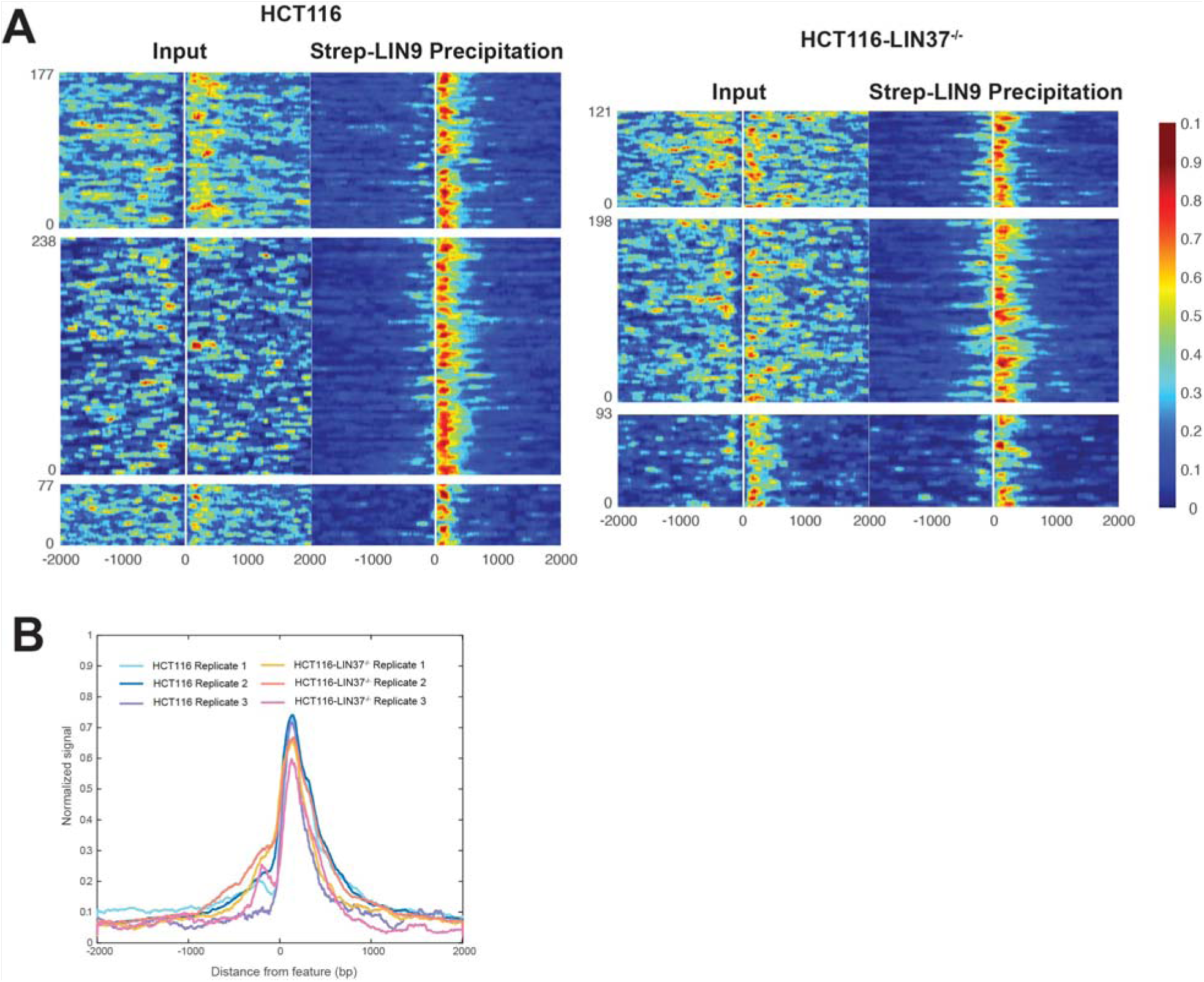
Comparison of MNase-ChIP replicate experiments in HCT116 and HCT116-LIN37^-/-^ cells. Replicates 1 and 3 are biological replicates. Replicate 2 is a technical replicate of Replicate 1 but using a 5-fold lower MNase concentration. (**A**) As in Figure 7B, heat maps for each experiment show the normalized DNA sequence read density across all DREAM genes that were detected with greater than 4.7-fold enrichment compared to a Strep-precipitation control. Replicates 1 to 3 are shown top to bottom in each cell type. (**B**) As in Figure 7B, overlay of the aggregated and normalized read signal density from the same set of genes as shown in the heat maps.

## Supplemental Methods

### DNA affinity purification

HCT116 cells were cultivated in 15cm plates and transfected with 70µl PEI and 15µg plasmids expressing wild-type and mutant Lin9 fused with an N-terminal 3xFlag tag. 24 hours after the transfection cells were treated with 5µM Nutlin-3a for 48 hours. Affinity purifications were performed as described earlier (Muller and Engeland, 2021). Biotinylated DNA probes were either amplified from the pGL4.10 empty vector or from pGL4.10 containing the mouse *Ccnb2* CDE/CHR MuvB-binding site (Muller et al., 2014). The following antibodies were applied for protein detection: FLAG-M2 (RRID:AB_262044, Sigma-Aldrich), p130/RBL2 (D9T7M) (RRID:AB_2798274, Cell Signaling), LIN37-T3 (custom-made at Pineda Antikörper-Service, Berlin, Germany) (Muller et al., 2017), Histone H3 (RRID:AB_331563, Cell Signaling Technology).

